# Independent derivations of the axis of arrhythmia for predicting drug-induced torsades de pointes

**DOI:** 10.1101/2024.09.18.613631

**Authors:** Stewart Heitmann, Jamie I Vandenberg, Adam P Hill

## Abstract

Torsades de pointes is a potentially lethal ventricular arrhythmia that may be caused by many classes of drugs. The likelihood of arrhythmia depends upon the particular combination of cardiac ion-channels that the drug targets. The risk for a given drug can be measured against the axis of arrhythmia, which is a conceptual line in the parameter space of ventricular cardiomyocyte electrophysiology. The axis describes the most potent combination of ion-channel blocks that is theoretically possible. It thus serves as a convenient yardstick for quantifying the pro-arrhythmic risk of novel drugs from their channel-blocking profiles. To date, the axis of arrhythmia has only been derived from biophysical computer simulations. Here, we derive the axis directly from the response profiles of selected ion-channels to known drugs (n=109). The new method provides an independent line of evidence for estimating the axis of arrhythmia. Using step-wise reduction, the two methods were found to converge to identical estimates of the axis in two ion-currents **(*I*_CaL_** and ***I*_Kr_)**. The resulting axis predicted the pro-arrhythmic risk of the test drugs with 89.9% to 91.7% accuracy, depending on the dose at which the drug was assessed. The axis of arrhythmia offers a practical method for predicting the pro-arrhythmic risk of novel drugs without the need for drug-specific computer simulations. It is the only metric of pro-arrhythmic risk that has been derived from both a biophysical model and a purely statistical model. It thus combines the benefits of biophysical interpretation and computational efficiency in assessing the torsadogenic risk of new drugs prior to clinical trials.

**Significance Statement:** Many classes of drugs can cause lethal cardiac arrhythmias by blocking ion-channels that are needed by heart cells for normal rhythm. International safety guidelines thus require all new drugs be tested in animal hearts or cell cultures prior to undertaking clinical trials in humans. Computer models offer an ethical alternative for drug testing that is gaining acceptance in the safety pharmacology community. However, the current corpus of models are usually too complex to apply outside of specialist computing laboratories. We recently proposed a new approach that eliminates the need for drug-specific computer simulations by using a large one-off computer simulation to identify the most pro-arrhythmic drug that is theoretically possible. Thereafter, that hypothetical drug can be used to assess the pro-arrhythmic risk of real drugs using only pen-and-paper calculations. In that study, the hypothetical drug was described by the *axis of arrhythmia* in the cardiac cell model. In the present study, we propose an alternative method for deriving the axis of arrhythmia directly from drug datasets. The two derivations are independent, yet they arrive at the same result. Together, they provide converging lines of evidence in support of this new approach to drug testing in safety pharmacology.

## Introduction

Pharmacological compounds can have off-target effects that disrupt the electrical activity of heart cells, potentially causing lethal cardiac arrhythmias such as torsades de pointes (Yap and Camm, 2003). The majority of torsadogenic drugs block the human ether-a-go-go related gene (hERG) channel which carries the major repolarizing current (*I_Kr_*) in ventricular cardiomyocytes (Perrin et al., 2008). Blocking hERG slows repolarization and consequently prolongs the ventricular action potential. The macroscopic effect is observed as prolongation of the QT interval of the electrocardiogram (Witchel, 2011). Prolongation is associated with an increased risk of spontaneous early after-depolarizations which can initiate torsades de pointes (Krummen et al., 2016). International safety guidelines (ICH, 2005) therefore recommend that new drugs be assessed for hERG block in vitro prior to conducting human trials.

Although all pro-arrhythmic drugs are known to block hERG, it does not follow that all drugs that block hERG are necessarily pro-arrhythmic (Martin et al., 2004; Hoffmann and Warner, 2006). For example, verapamil is an approved anti-arrhythmic drug that simultaneously blocks the hERG outward current and the L-type calcium inward current (Singh et al., 1978). It is not pro-arrhythmic because the inward and outward currents remain balanced, nevertheless it would fail the standard hERG test. Indeed, it is estimated that up to 70% of drugs block hERG (Shah, 2005), yet only 4% cause arrhythmia (Aiba et al., 2005). To rectify this inefficiency, the safety pharmacology community are actively pursuing more accurate safety assays that target multiple ion-channels (Pugsley et al., 2008; Colatsky et al., 2016; Strauss et al., 2021), including *I*_CaL_, *I*_Kr_, *I*_NaL_ and *I*_Ks_. Computational models of the effects of drugs on cardiac electrophysiology are an important part of that effort, although biophysically detailed models require substantial computing resources and expertise which often hinder their widespread adoption (Heitmann et al., 2023).

### Axis of Arrhythmia

To eliminate the technological barrier posed by drug-specific computer simulations, we proposed the concept of the *axis of arrhythmia* (Heitmann et al., 2023) for comparing the multi-channel blocking signature of a drug to the most pro-arrhythmic combination of ion-channel blocks that is theoretically possible. In that study, the axis was derived from simulated ventricular cardiomyocytes with randomly expressed ion-channels. Cardiomyocytes that elicited early after-depolarizations were classified as ectopic (pro-arrhythmic) and those that did not were classified as benign (Figure 1A). The two classes of cardiomyocytes occupy distinct regions of parameter space that were readily separated by a linear classification boundary (Figure 1B). The axis of arrhythmia (solid line) describes the optimal direction for shifting a cardiomyocyte from the benign parameter regime (white region) towards the ectopic parameter regime (red region) by altering the conductivity of the ion-channels (*G*_ion_).

**Figure 1.**
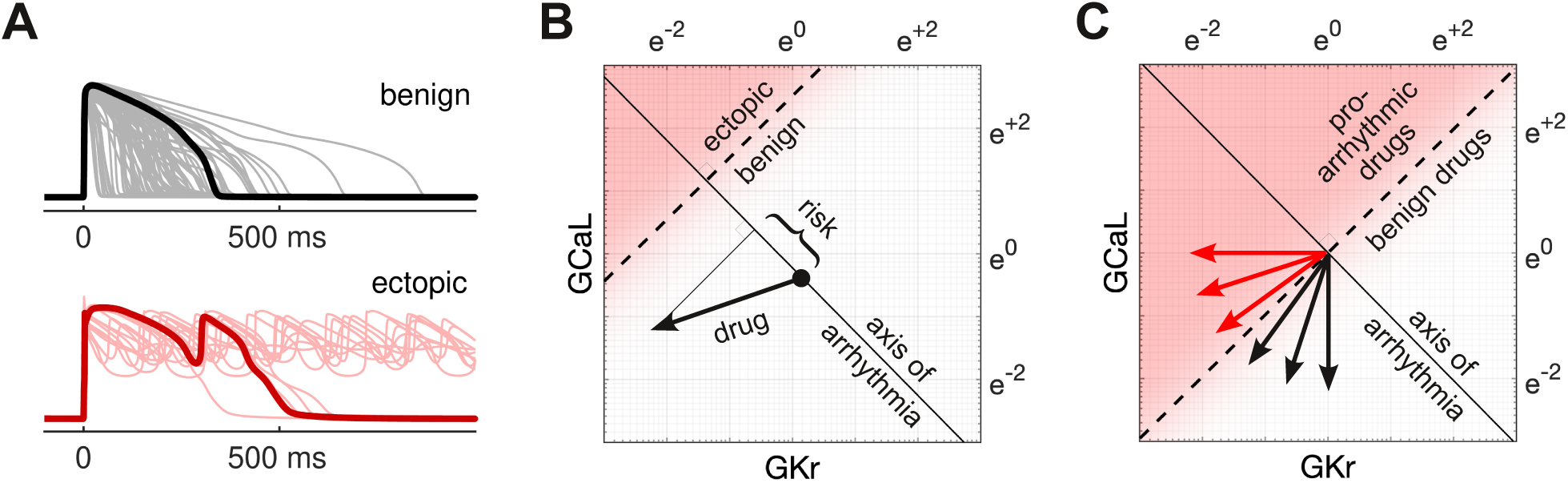
The axis of arrhythmia. (**A**) Simulated action potentials for a large population of ventricular cardiomyocytes with randomized ion-channel conductivity. Cardiomyocytes that exhibited early after-depolarizations (red) were classified as ectopic and the remainder were classified as benign. (**B**) Parameter map showing the relationship between ectopic (red region) and benign (white region) cardiomyocytes and the conductivity of two ion-channels (*G_Kr_* and *G_CaL_*) in logarithmic coordinates. The axis of arrhythmia (solid line) is orthogonal to the classification boundary (dashed) by definition. It describes the optimal path for transforming a benign cardiomyocte into an ectopic cardiomyocyte. The axis serves as a yardstick for measuring the pro-arrhythmic risk of a channel-blocking drug (arrow) without the need for drug-specific computer simulations. (**C**) The proposed method for deriving the axis of arrhythmia directly from the channel-blocking characteristics of known drugs. In this approach, the axis of arrhythmia is derived from the boundary (dashed) that optimally separates pro-arrhythmic drugs (red arrows) from benign drugs (black arrows). Panels A and B are reproduced from Heitmann et al. (2023).

### Risk Metric

The pro-arrhythmic component of a drug is calculated by projecting the drug’s action (Figure 1B, arrow) onto the axis of arrhythmia using vector geometry (Heitmann et al., 2023). Specifically, the pro-arrhythmic risk score,

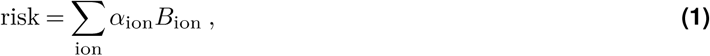

is the length of that projection, where *α*_ion_ is the effect of the drug on the ion-current, and *B*_ion_ is the corresponding coefficient of the axis of arrhythmia. Using four ion-currents (*I*_CaL_, *I*_Kr_, *I*_NaL_, *I*_Ks_), Eq. (1) predicted the clinical risk of 109 drugs with an accuracy of 88.1% to 90.8%, depending on the dose at which the drug was assessed (Heitmann et al., 2023).

### Complimentary Derivations

The present study provides a new method for deriving the axis of arrhythmia, in this case from the potency data of drugs with known clinical risk labels. This new method uses statistical analysis to identify the linear boundary that separates the trajectories of pro-arrhythmic versus benign drugs (Figure 1C). The axis of arrhythmia is then inferred from the orientation of the classification boundary. Through a process of step-wise reduction, we will show that the two independent derivations converge to the same result. Namely, a metric that uses the axis of arrhythmia in two ion-currents (*I*_CaL_ and *I*_Kr_) to predict the pro-arrhythmic risk of drugs with an accuracy of 90.8% to 91.7%.

## Results

The Hill equation (Hill, 1910) was used to reconstruct the dose responses of four ion-currents (*I*_CaL_, *I*_Kr_, *I*_NaL_, *I*_Ks_) from the half-maximal inhibitory concentrations (*IC*_50_) of drugs with known clinical risk labels. In this case, the drugs (n=109) were from a publicly available dataset curated by Llopis-Lorente et al. (2020). The conductivity of the drugged ion-channel was expressed as a dose-dependent fraction (0 *≤ δ ≤* 1) of its baseline conductance. Specifically,

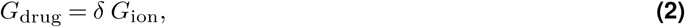

where *δ* = 0 corresponds to 100% block of the channel, and *δ* = 1 corresponds to 0% block (Figure 2A). Following Heitmann et al. (2023), the drug was analyzed independently of the cardiomyocytes by using the logarithmic transform to convert Eq. (2) into an additive expression,

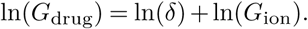

**Figure 2.**
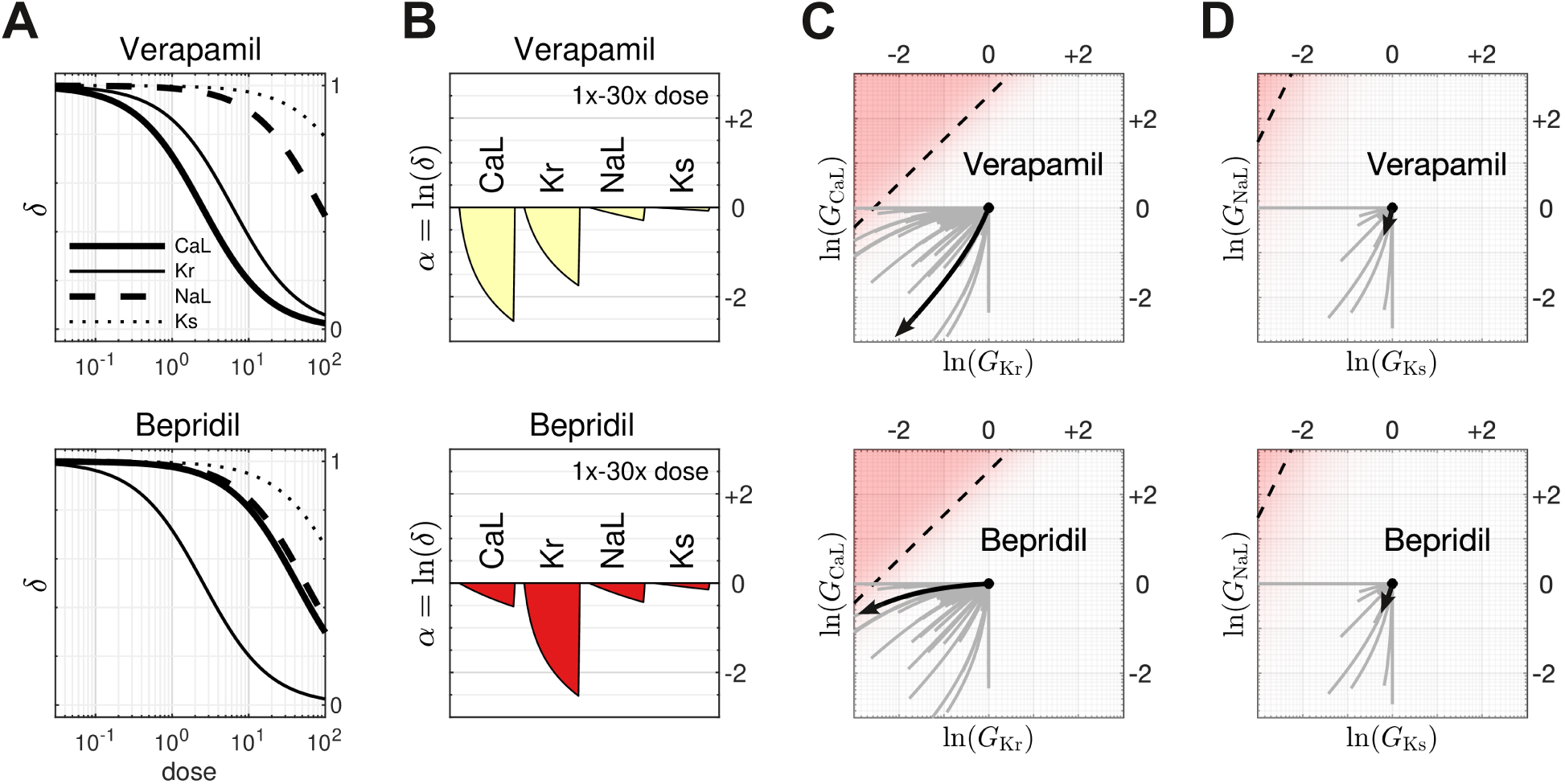
Examples of multi-channel drug-block. Verapamil is an anti-arrhythmic drug that primarily blocks *I*_CaL_ and *I*_Kr_. Bepridil is a pro-arrhythmic drug that primarily blocks *I_Kr_*. (**A**) Dose-response profiles of *I_CaL_*, *I_Kr_*, *I_NaL_* and *I_Ks_* to verapamil (top) and bepridil (bottom). The doses are normalized to the therapeutic dose. (**B**) The dose-response profiles in logarithmic coordinates (*α* = ln *δ*), shown side-by-side. Each profile spans 1x to 30x therapeutic dose. (**C**) The corresponding dose-response trajectories (black) in *G*_Kr_ versus *G*_CaL_. The trajectories of all other drugs are shown in gray. The ectopic parameter regime (shaded red) of the simulated cardiomyocytes in Heitmann et al. (2023) is included for comparison. (**D**) Case of *G*_Ks_ versus *G*_NaL_.

The action of the drug in the logarithmic coordinate frame is written as *α* = ln(*δ*). It describes the shift in the conductance of the channel caused by a given dose of drug (Figure 2B). Importantly, *α* is independent of the baseline conductance (*G*_ion_) of the cell phenotype. For the case of four ion-channels, the simultaneous action of the drug is described by the vector,

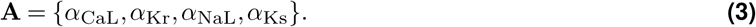

It traces out a curvilinear trajectory with dose (Figures 2C,D) because each ion-channel responds to the drug at a differing rate. We used statistical analysis to find the linear boundary that optimally segregated the trajectories of pro-arrhythmic drugs from the trajectories of benign drugs.

### Deriving the axis of arrhythmia from the drug trajectories

In the context of a cardiomyocyte, the axis of arrhythmia describes the optimal combination of channel blocks for inducing early after-depolarizations in the ventricular action potential (Heitmann et al., 2023). Here, we derive the axis of arrhythmia directly from the drug trajectories without resort to simulating the action potential. The drugs were segregated into two classes (benign versus pro-arrhythmic) according to their clinical labels from the Credible Meds consortium (Methods). The linear boundary that optimally separated those classes (dashed line in Figures 3A,B) was obtained by multivariate logistic regression,

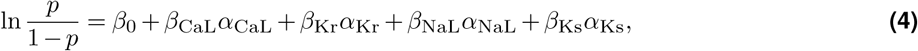

which predicts the log-odds of a drug being pro-arrhythmic, where *{α*_CaL_, *α*_Kr_, *α*_NaL_, *α*_Ks_*}* are the predictor variables, and *{β*_CaL_, *β*_Kr_, *β*_NaL_, *β*_Ks_*}* are the regression coefficients to be estimated. The classification boundary corresponds to the four-dimensional plane where the probability of the drug being pro-arrhythmic is exactly *p* = 0.5. Since we know *a priori* that zero dose of drug imposes zero channel block, the boundary was pinned to the origin by imposing *β*_0_ = 0.

**Figure 3.**
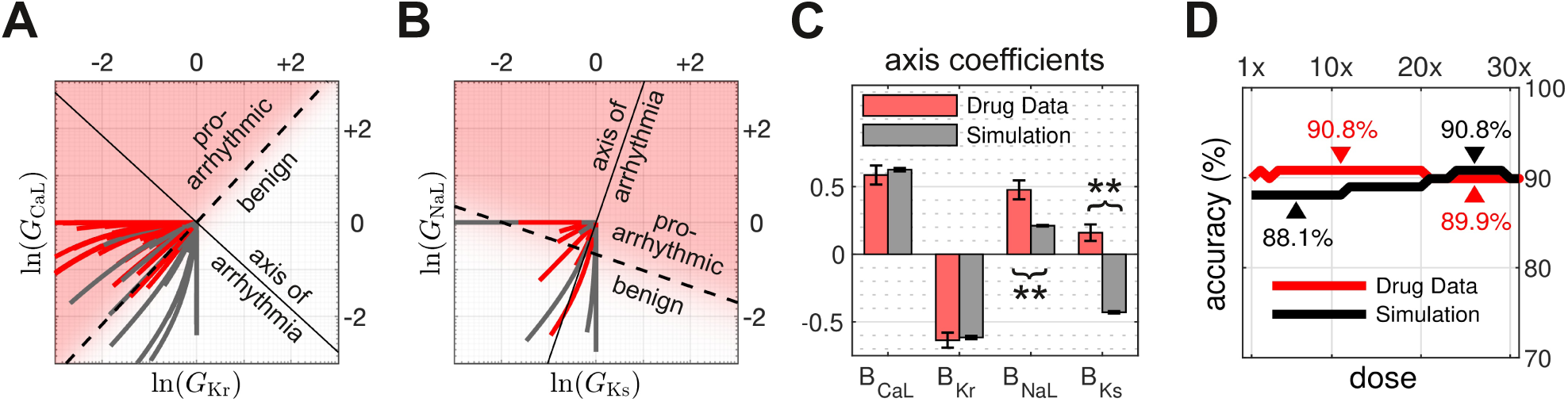
Analyzing drug-block in four ion-currents. (**A**) The axis of arrhythmia (solid line) that was derived from the linear classification boundary (dashed line) that separates the pro-arrhythmic drugs (red) from the benign drugs (gray) in the four-dimensional space of ionic conductances (*G*_CaL_*, G*_Kr_*, G*_NaL_*, G*_Ks_). The trajectories of drug-block are visualized by projecting them onto the two-dimensional plane where ln(*G*_Ks_) = 0 and ln(*G*_NaL_) = 0. (**B**) Same as panel A, for the two-dimensional plane where ln(*G*_Kr_) = *−*0.5 and ln(*G*_CaL_) = 0. (**C**) Coefficients of the axis of arrhythmia. Red bars are the estimates obtained from the drug trajectories. Gray bars are the estimates from the cardiac cell simulations in Heitmann et al. (2023). Error bars are the 95% confidence intervals. Significant differences (*p <* 0.001) are marked with a double asterisk. (**D**) Accuracy profiles of the predictions of pro-arrhythmic risk made by the drug-derived axis of arrhythmia (red) versus the simulation-derived axis (black). The simulation results are reproduced from Figure 8A of Heitmann et al. (2023).

By definition, the regression coefficients *{β*_CaL_, *β*_Kr_, *β*_NaL_, *β*_Ks_*}* constitute a vector that is orthogonal to the classification boundary. As such, the coefficients define the basis vector of the axis of arrhythmia,

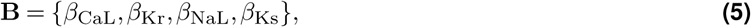

which was normalized to unit length, *||***B***||* = 1.

### Coefficients of the axis of arrhythmia

Leave-one-out cross-validation was used to train the regression on all combinations of *n−*1 training drugs, while setting aside one drug from each block for testing. The estimated coefficients were re-scaled to unit length, and averaged across all blocks. The averaged coefficients *{B*_CaL_, *B*_Kr_, *B*_NaL_, *B*_Ks_*}* are listed in Table 1 (Drug Dataset) and plotted in Figure 3C (red). All four ion-channels were found to be statistically significant predictors of the clinical risk labels of the drugs (*p <* 0.001).

**Table 1.** Coefficients of the axis of arrhythmia in four ion-currents. Two sets of coefficients *{B*_CaL_*, B*_Kr_*, B*_NaL_*, B*_Ks_*}* were estimated independently from the drug dataset and the cardiac cell simulations. The drug-based estimates are the averages of 108 leave-one-out training blocks with 32 doses per drug. The simulation-based estimates are reproduced from Heitmann et al. (2023). Confidence intervals are 95% of the t-distribution. Welch’s two-sample t-test is used to compare the estimates from the drug dataset with those from the cell simulation. Statistically significant p-values (*p <* 0.001) are marked with a double asterisk.

| Coefficient | Drug Dataset<br>($n=108 \times 32$) | Cell Simulation<br>($n=100,000$) | Two-Sample<br>Welch's t-test |
| --- | --- | --- | --- |
| $B_{CaL}$ | $0.586 \pm 0.070$<br>$SE = 0.036$<br>$t(3452) = 16.4$<br>$p < 0.001^{**}$ | $0.625 \pm 0.012$<br>$SE = 0.0061$<br>$t(99995) = 102.8$<br>$p < 0.001^{**}$ | $t(3659) = -1.09$<br>$p = 0.28$ |
| $B_{Kr}$ | $-0.636 \pm 0.056$<br>$SE = 0.028$<br>$t(3452) = -22.4$<br>$p < 0.001^{**}$ | $-0.616 \pm 0.012$<br>$SE = 0.0060$<br>$t(99995) = -102.4$<br>$p < 0.001^{**}$ | $t(3771) = -0.69$<br>$p = 0.49$ |
| $B_{NaL}$ | $0.476 \pm 0.070$<br>$SE = 0.036$<br>$t(3452) = 13.3$<br>$p < 0.001^{**}$ | $0.211 \pm 0.006$<br>$SE = 0.0030$<br>$t(99995) = 69.9$<br>$p < 0.001^{**}$ | $t(3504) = 7.36$<br>$p < 0.001^{**}$ |
| $B_{Ks}$ | $0.160 \pm 0.061$<br>$SE = 0.031$<br>$t(3452) = 5.11$<br>$p < 0.001^{**}$ | $-0.430 \pm 0.009$<br>$SE = 0.0045$<br>$t(99995) = -94.7$<br>$p < 0.001^{**}$ | $t(3602) = 18.7$<br>$p < 0.001^{**}$ |
| Model | $R^2 = 0.43$ | $R^2 = 0.76$ | |

For comparison, the corresponding coefficients from the cardiac cell simulations (Heitmann et al., 2023) are reproduced in Table 1 (Cell Simulation) and plotted in Figure 3C (gray). Welch’s two-sample t-test confirmed that the competing estimates of *B*_CaL_ were virtually identical, *t*(3659) = *−*1.09, *p* = 0.28, as were the competing estimates of *B*_Kr_, *t*(3771) = *−*0.69, *p* = 0.49. Nevertheless, the two methods did not agree on *B*_NaL_ which had significantly different estimates, *t*(3504) = 7.36, *p <* 0.001. Nor did the methods agree on *B*_Ks_ which were likewise significantly different, *t*(3602) = 18.7, *p <* 0.001, and had opposite sign.

### Prediction accuracy

Despite the discrepancies in the coefficients from the drug dataset versus the cell simulation, the four ion-channels were all statistically significant predictors in both models (*p <* 0.001). The drug-based model predicted the risk labels of the test drugs with 89.9% to 90.8% accuracy, depending on the dose at which the drug was assessed (Figure 3D, red). Similarly, the simulation-based model (black) predicted the risk labels with 88.1% to 90.8% accuracy (reproduced from Heitmann et al., 2023). In this regard, the two methods gave essentially the same result. We concluded that the discrepancies in the coefficients must be reflecting underlying discrepancies in the drug data and the cell model, each of which have recognized limitations (Kramer et al., 2020; Colatsky et al., 2016). Since it is unknown which method is more correct, we undertook stepwise elimination of those predictors that were inconsistent across both methods, beginning with *B*_Ks_.

### The axis of arrhythmia in three ion-currents

The analysis was repeated using three conductance parameters only (*G*_CaL_*, G*_Kr_*, G*_NaL_). As before, the axis of arrhythmia was estimated from the classification boundary that optimally separated the trajectories of pro-arrhythmic versus benign drugs in the dataset (Figure 4A). New cell simulations were also conducted using random *G*_CaL_*, G*_Kr_*, G*_NaL_ only, whereupon the axis of arrhythmia was estimated from the classification boundary that separated the ectopic cell phenotypes from the benign cell phenotypes (Figure 4B).

**Figure 4.**
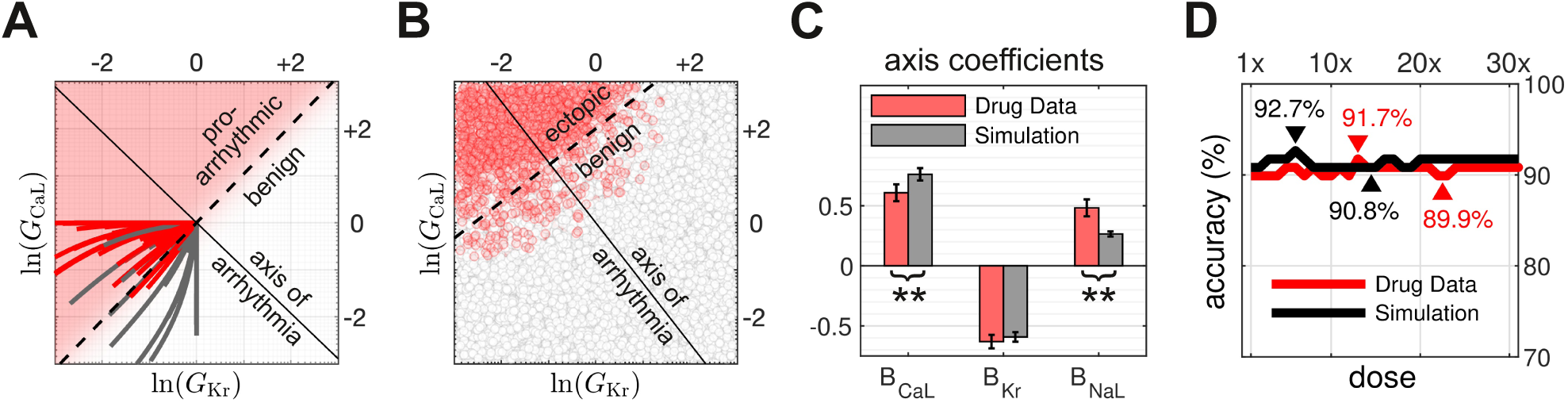
Analyzing drug-block in three ion-currents. (**A**) The axis of arrhythmia (solid line) from the trajectories of drug-block in three conductance parameters (*G*_CaL_*, G*_Kr_*, G*_NaL_). The trajectories are visualized by projecting them onto the two-dimensional plane where ln(*G*_NaL_) = 0. The dashed line indicates where the three-dimensional classification boundary intersects the two-dimensional plane. (**B**) The corresponding axis of arrhythmia from the simulated cardiac cells (n=10000). The parameters (*G*_CaL_*, G*_Kr_*, G*_NaL_) of each cell were selected at random. Each cell was classified as either benign (white) or ectopic (red) using early after-depolarizations as the biomarker. The axis of arrhythmia was obtained from the classification boundary that optimally separated the benign and ectopic cell phenotypes. (**C**) Coefficients of the axis of arrhythmia. Red bars are the estimates from the drug trajectories. Gray bars are the estimates from the cell simulations. Error bars are the 95% confidence intervals. Significant differences (*p <* 0.001) are marked with a double asterisk. (**D**) Accuracy of the predictions of pro-arrhythmic risk made by the axis of arrhythmia from the drug trajectories (red) versus the axis of arrhythmia from the cell simulations (black).

The resulting estimates of the coefficients of the axis of arrhythmia *{B*_CaL_, *B*_Kr_, *B*_NaL_*}* are listed in Table 2. All three were found to be significant predictors (*p <* 0.001) in both methods. However the estimates from the drug trajectories were only in partial agreement with the estimates from the cell simulations (Figure 4C, red versus gray). Welch’s two-sample t-test confirmed that the competing estimates of *B*_Kr_ were statistically equivalent, *t*(7182) = *−*1.12, *p* = 0.26, but the estimates of *B*_CaL_ and *B*_NaL_ were statistically different, *t*(7406) = *−*3.37, *p <* 0.001 and *t*(4089) = 5.76, *p <* 0.001, respectively. Despite these differences, the two models again had near-identical accuracy profiles (Figure 4D). Depending on the dose at which the drugs were assessed, the drug-based model (red) predicted the clinical risk labels with 89.9% to 91.7% accuracy, and the simulation-based model (black) predicted them with 90.8% to 92.7% accuracy.

**Table 2.** Coefficients of the axis of arrhythmia in three ion-currents. The coefficients were estimated independently from the drug dataset and simulated cardiac action potentials. Confidence intervals are 95% of the t-distribution. Statistically significant differences (*p <* 0.001) are marked with a double asterisk.

| Coefficient | Drug Dataset<br>($n=108 \times 32$) | Cell Simulation<br>($n=10,000$) | Two-Sample<br>Welch's t-test |
| --- | --- | --- | --- |
| $B_{CaL}$ | $0.608 \pm 0.070$<br>$SE = 0.036$<br>$t(3453) = 17.0$<br>$p < 0.001$ ** | $0.762 \pm 0.051$<br>$SE = 0.026$<br>$t(9996) = 29.2$<br>$p < 0.001$ ** | $t(7406) = -3.47$<br>$p < 0.001$ ** |
| $B_{Kr}$ | $-0.631 \pm 0.056$<br>$SE = 0.029$<br>$t(3453) = -22.1$<br>$p < 0.001$ ** | $-0.591 \pm 0.040$<br>$SE = 0.020$<br>$t(9996) = -29.2$<br>$p < 0.001$ ** | $t(7182) = -1.12$<br>$p = 0.26$ |
| $B_{NaL}$ | $0.482 \pm 0.071$<br>$SE = 0.036$<br>$t(3453) = 13.3$<br>$p < 0.001$ ** | $0.265 \pm 0.021$<br>$SE = 0.011$<br>$t(9996) = 24.5$<br>$p < 0.001$ ** | $t(4089) = 5.76$<br>$p < 0.001$ ** |
| Model | $R^2 = 0.41$ | $R^2 = 0.82$ | |

### The axis of arrhythmia in two ion-currents

Having found that the drug data and the simulations disagreed on the estimates of *B*_NaL_, we eliminated *I*_NaL_ from further consideration. The axis of arrhythmia in the two remaining ion-currents (*I*_CaL_ and *I*_Kr_) was then derived from the trajectories of drug-block in *G*_CaL_ versus *G*_Kr_ (Figure 5A) and also from a fresh set of cardiac cell simulations where *G*_CaL_ and *G*_Kr_ were the only parameters manipulated (Figure 5B). The resulting estimates of the coefficients *{B*_CaL_, *B*_Kr_*}* are listed in Table 3 and plotted in Figure 5C. The two ion-currents were once again found to be significant predictors in both models (*p <* 0.001), but this time the respective estimates of *B*_CaL_ and *B*_Kr_ were each found to be statistically equivalent, *t*(2941) = *−*0.80, *p* = 0.42 and *t*(3093) = *−*1.68, *p* = 0.093, meaning that the two models returned the same result. Both models also had virtually identical accuracy profiles (Figure 5C) wherein the drug-based model (red) predicted the clinical risk labels of the drugs with 89.0% to 90.8% accuracy, and the simulation-based model (black) predicted the risk labels with 89.9 % to 91.7% accuracy, depending on dose. Based on the equivalence of the coefficients of the axes, and the equivalence of the accuracy profiles, we concluded that the independent derivations of the axis of arrhythmia had converged to the same findings.

**Figure 5.**
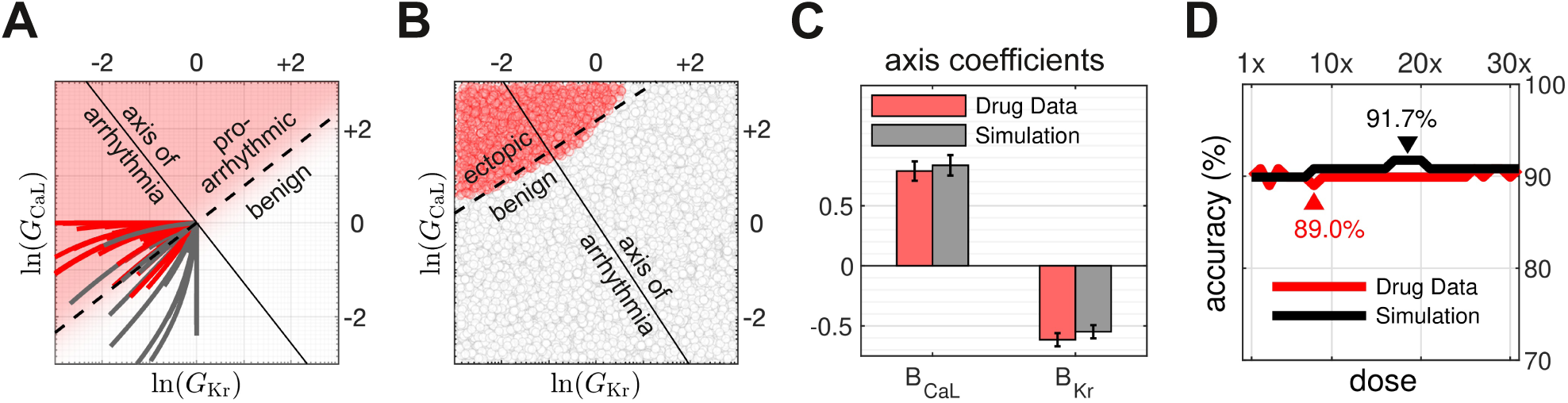
Analyzing drug-block in two ion-currents. (**A**) The axis of arrhythmia from the trajectories of drug-block in two conductance parameters (*G*_CaL_*, G*_Kr_). (**B**) The corresponding axis of arrhythmia from the simulated cardiac cells (n=10000). (**C**) The coefficients of the axis of arrhythmia. Red bars are the estimates from the drug trajectories. Gray bars are the estimates from the cell simulations. Error bars are the 95% confidence intervals. No statistical differences were observed in the estimates. (**D**) Accuracy of the pro-arrhythmic risk predictions made by the axis of arrhythmia from the drug trajectories (red) versus that from the cell simulations (black).

**Table 3.** Coefficients of the axis of arrhythmia in two ion-currents. The coefficients *B*_CaL_ and *B*_Kr_ were estimated both from the trajectories of drug-block in *G*_CaL_ and *G*_Kr_ and also from the phenotypes of simulated cardiac cells in which *G*_CaL_ and *G*_Kr_ were randomly manipulated. Confidence intervals are 95%. Significant differences (*p <* 0.001) are marked with a double asterisk.

| Coefficient | Drug Dataset<br>($n=108 \times 32$) | Cell Simulation<br>($n=10,000$) | Two-Sample<br>Welch's t-test |
| --- | --- | --- | --- |
| $B_{CaL}$ | $0.789 \pm 0.081$<br>$SE = 0.041$<br>$t(3454) = 19.1$<br>$p < 0.001^{**}$ | $0.836 \pm 0.085$<br>$SE = 0.043$<br>$t(9997) = 19.4$<br>$p < 0.001^{**}$ | $t(2941) = -0.80$<br>$p = 0.42$ |
| $B_{Kr}$ | $-0.615 \pm 0.055$<br>$SE = 0.028$<br>$t(3454) = -22.0$<br>$p < 0.001^{**}$ | $-0.548 \pm 0.055$<br>$SE = 0.028$<br>$t(9997) = -19.5$<br>$p < 0.001^{**}$ | $t(3093) = -1.68$<br>$p = 0.093$ |
| Model | $R^2 = 0.37$ | $R^2 = 0.91$ | |

### Pooled axis of arrhythmia

Having concluded that the competing estimates of the axes of arrhythmia in two ion-currents were statistically equivalent (Table 3), we averaged them to obtain the pooled estimate of the axis (Figure 6A). It represents the combined estimate of the axis from both lines of evidence, where *B*_CaL_ = 0.813 *±* 0.12 and *B*_Kr_ = *−*0.582 *±* 0.08 (Table 4; left column). These are the values that we recommend using with Eq. (1) to best predict the pro-arrhythmic risk of a drug from *I*_CaL_ and *I*_Kr_ block. The accuracy of these coefficients in predicting the clinical labels (benign versus pro-arrhythmic) of the test drugs ranged from 89.9% to 91.7% (Figure 6B, black). The scoring thresholds (Figure 6B, red) were optimized to achieve the highest possible classification accuracy at each dose. Those thresholds can be tuned to achieve the desired balance between false-positives and true-positives in the Receiver Operating Characteristics of the metric (Figure 6C). Alternatively, the raw score can be used to judge the pro-arrhythmic risk of a drug without necessarily converting it into a binary classification. It describes how far the drug shifts the electrophysiology toward (or away) from the pro-arrhythmic region of parameter space.

**Figure 6.**
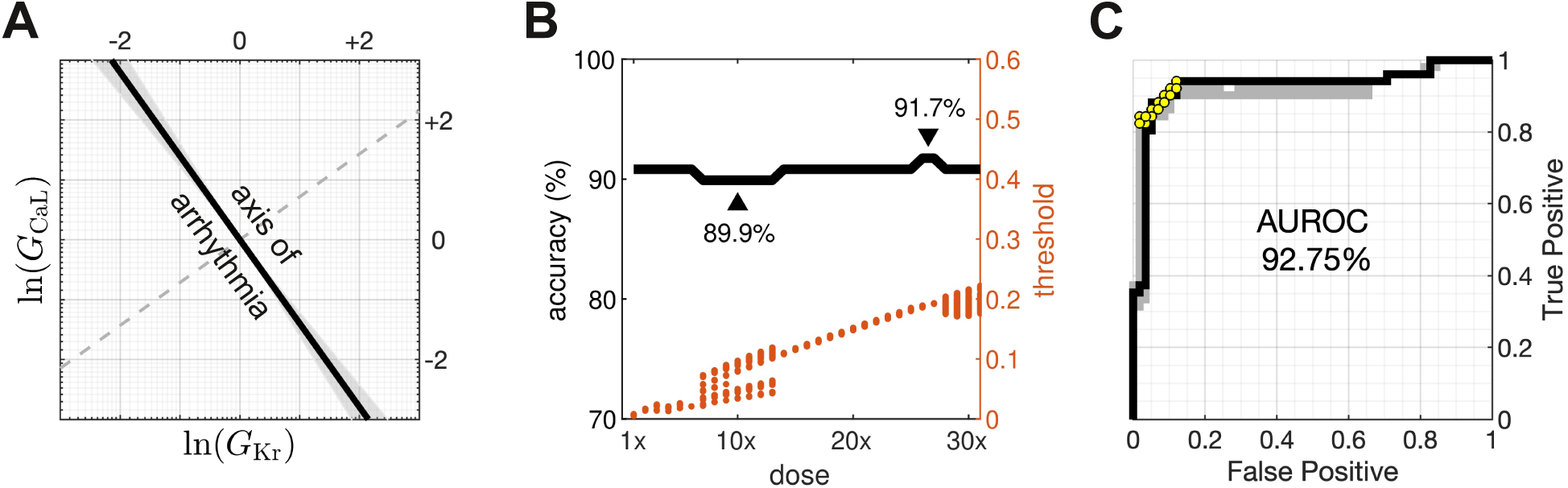
Pooled axis of arrhythmia in two ion-currents. (**A**) The pooled axis (black line) was obtained by averaging the coefficients from the drug dataset and the cardiac cell simulations. Shaded area is the 95% confidence envelope. (**B**) The accuracy profile (black line) of the predictions of pro-arrhythmic risk by the pooled axis of arrhythmia. The optimal scoring thresholds for each dose are shown in red. Multiple scoring thresholds are optimal for some doses. (**C**) Receiver Operating Characteristics (ROC) of the risk metric for drugs tested at 3x therapeutic dose. AUROC is the area under the ROC curve. The ROC curves for all other doses are shown in gray. Yellow dots correspond to the accuracy rates in panel B.

**Table 4.** Pooled coefficients of the axis of arrhythmia. The pooled estimates of the coefficients in two ion-currents (*I*_CaL_*, I*_Kr_) and three ion-currents (*I*_CaL_*, I*_Kr_*, I*_NaL_). Confidence intervals are 95% (*±*1.96 SE).

| Coefficient | Two Currents | Three Currents |
| --- | --- | --- |
| $B_{CaL}$ | $0.813 \pm 0.12$<br>$SE = 0.060$ | $0.691 \pm 0.087$<br>$SE = 0.045$ |
| $B_{Kr}$ | $-0.582 \pm 0.078$<br>$SE = 0.040$ | $-0.617 \pm 0.069$<br>$SE = 0.035$ |
| $B_{NaL}$ | – | $0.377 \pm 0.075$<br>$SE = 0.038$ |

### Drug risk profiles

The computed risk scores for eight selected drugs are shown in Figure 7. Each panel represents one drug, where the heavy black line indicates the risk scores for a range of doses, up to 30x the therapeutic dose. Drugs that scored above the threshold (pink region) were predicted to be pro-arrhythmic, and those that did not were predicted to be benign. The torsadogenic risk labels from Credible Meds served as the ground truth. Of the selected drugs that were correctly classified (Figure 7A), the scores for bepridil (class 1) and dolasetron (class 2) were unambiguously *above* the threshold (pro-arrhythmic). Likewise, the scores for quinine (class 3) and verapamil (class 4) were unambiguously *below* the threshold (benign). Of the selected drugs that were incorrectly classified (Figure 7B), the computed scores yielded false negatives for sotalol (class 1) and saquinavir (class 2), as well as false positives for ranolazine (class 3) and propafenone (class 3). Ranolazine was the only drug whose classification was rescued by the use of three ion-currents (dotted line). It substantially blocks *I*_NaL_ in addition to *I*_CaL_ and *I*_Kr_ while preserving the overall balance of inward and outward currents. Propafenone is an example of a borderline case where the scores straddle the threshold, producing false-positive classifications at low doses and true-negatives at high doses. Approximately half of all the incorrectly classified drugs in the dataset were borderline cases, of which propafenone is perhaps the most extreme example.

**Figure 7.**
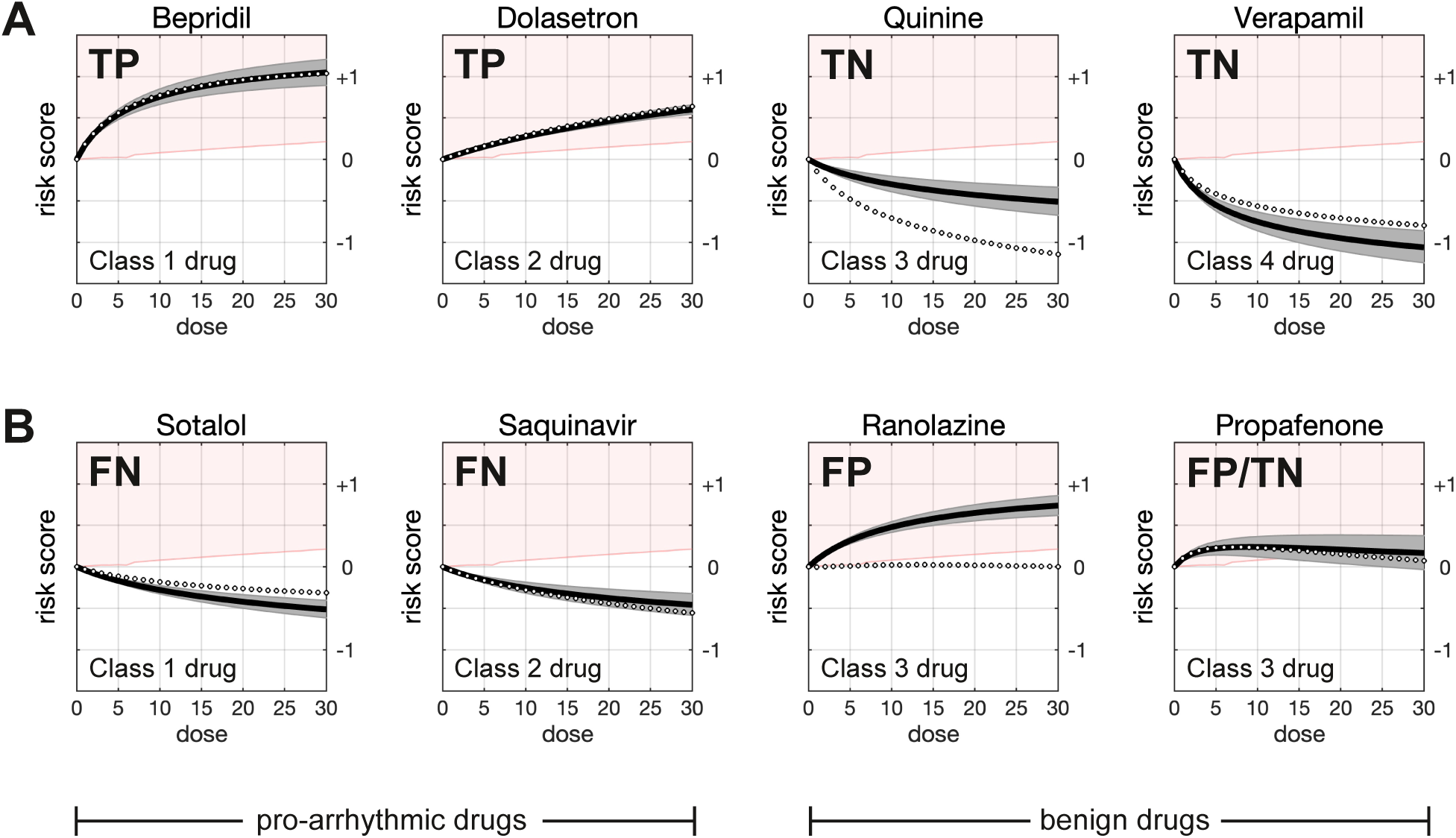
Pro-arrhythmic risk scores for selected drugs. Each panel shows the risk scores (heavy black line) for a range of doses of a given drug. The scores were calculated using the pooled axis of arrhythmia in two ion-currents (*β*_CaL_ = 0.813 *±* 0.12*, β*_Kr_ = *−*0.582 *±* 0.078) and include a 95% confidence interval (shaded gray). Drugs with scores in the pink region exceeded the scoring threshold and were classified as pro-arrhythmic accordingly. Scores in the white region were classified as benign. For comparison, the dotted line shows the risk scores that were calculated using the pooled axis in three ion-currents (*β*_CaL_ = 0.691 *±* 0.087*, β*_Kr_ = *−*0.617 *±* 0.069*, β*_NaL_ = 0.377 *±* 0.075). (**A**) Examples of drugs that were correctly classified by the risk metric. Each drug is drawn from one of the four torsadogenic risk classes from Credible Meds, where we regarded class 1 and 2 drugs as *pro-arrhythmic*, and class 3 and 4 drugs as *benign*. TP denotes true-positive. TN denotes true-negative. (**B**) Examples of drugs that were incorrectly classified. FN denotes false-negative. FP denotes false-positive. No class 4 drugs were misclassified. Ranolazine was the only drug in the dataset that was rescued by the use of three ion-currents (*I*_CaL_*, I*_Kr_*, I*_NaL_) instead of two (*I*_CaL_*, I*_Kr_). See the Supplementary Information for the risk profiles of all drugs in the dataset.

### Comparison to the hERG safety margin

The conventional hERG test (Webster et al., 2002; Redfern et al., 2003; Leishman et al., 2020) deems a drug to carry a liability for torsades de pointes if the margin between effective free therapeutic plasma concentration (EFTPC) and the *IC*_50_ of *I*_Kr_ is too small. A 30-fold margin is commonly used as the threshold for safety. The drugs in the present dataset were classified according to that criteria, after excluding Lidocaine and Rufinamide for missing hERG data. The 30-fold hERG safety margin was found to be 86.0% accurate in predicting the clinical risk labels (Figure 8A). In comparison, the pooled axis of arrhythmia in two ion-channels was 90.8% accurate (Figure 8B). In this case, the drugs were assessed at 4x therapeutic dose, which is similar to that used by the qNet metric of torsadogenic risk (Li et al., 2019) from the Comprehensive in Vitro Proarrhythmia Assay (CiPA) initiative (Colatsky et al., 2016).

**Figure 8.**
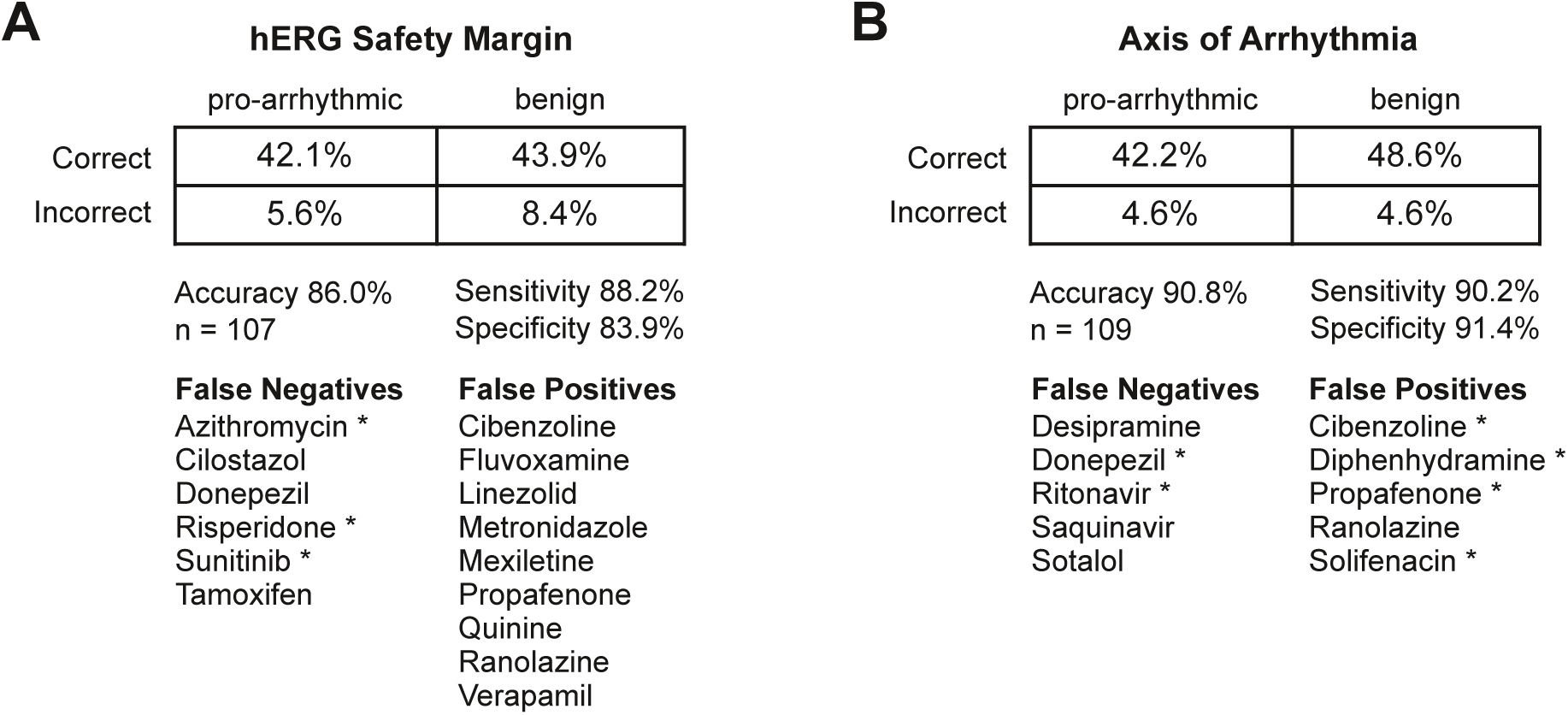
Accuracy of the conventional hERG assay compared to axis of arrhythmia. (**A**) The 2 x 2 contingency table for the conventional hERG assay using a 30-fold safety margin. The hERG assay was 86.0% accurate in predicting the clinical risk labels. Misclassified drugs are listed below the contingency table. Borderline cases are marked with an asterisk. Drugs with missing *IC*_50_ values for *I*_Kr_ were excluded from the analysis. (**B**) The 2 x 2 contingency table for the pooled axis of arrhythmia in two ion-channels. The drugs were assessed at 4x therapeutic dose. The axis of arrhythmia was 90.8% accurate in predicting the clinical risk labels. Approximately half of the misclassified drugs were borderline cases, as indicated by the asterisks.

## Discussion

The axis of arrhythmia (Heitmann et al., 2023) is a conceptual line in the parameter space of ventricular cardiomyocte electrophysiology. It describes the optimal combination of ion-channel blocks for shifting the electrophysiology toward the pro-arrhythmic regime where early after-depolarizations arise in the action potential. The axis serves as a convenient yardstick for quantifying the torsadogenic risk posed by drugs where pore-block is the primary mechanism of action. As such, it forms the basis of a simple metric for predicting the pro-arrhythmic risk of novel drugs prior to clinical trials. The metric improves upon standard hERG assay by integrating evidence from multiple ion-channels, notably *I*_Kr_ and *I*_CaL_. Under current ICH guidelines (ICH, 2005) the metric would only be applied in cases where follow-up studies to the core battery (*in vitro I*_Kr_ and *in vivo* QT) are required (Strauss et al., 2021). Although the metric could potentially be included in the core assay itself because it only requires one additional current (*I*_CaL_).

### Independent lines of evidence

The axis of arrhythmia was originally derived from biophysical computer simulations where the ion-channels were manipulated at random (Heitmann et al., 2023). The present study offers a new and independent method for deriving the axis of arrhythmia directly from the multi-channel potencies of drugs with known clinical risk labels. Together, the two methods provide converging lines of evidence for the pro-arrhythmic effects of multi-channel drug-block in ventricular cardiomyocytes. The pooled results from both methods give the best estimate of the true axis of arrhythmia. We know of no other model of pro-arrhythmic risk that can be derived independently from both a biophysical model and a statistical model.

### Convergent findings

The original formulation of the axis used four ion-currents (*I*_CaL_*, I*_Kr_*, I*_NaL_*, I*_Ks_), all of which were significant predictors of early after-depolarizations (Heitmann et al., 2023). The present study found that those same four currents were also significant predictors of the clinical labels of torsadogenic risk in the drug dataset. Despite their differing origins, the two methods were in near-perfect agreement for the contributions of *I*_CaL_ and *I*_Kr_, although they disagreed on the contributions of *I*_NaL_ and *I*_Ks_. Those discrepancies seemed to be at odds with the findings that (i) all four parameters were significant predictors in their own right, and (ii) the axes obtained from both methods were equally accurate at predicting the pro-arrhythmic risk of the test drugs. We concluded that the discrepant coefficients must have been compensating for inconsistencies between the biophysical model, the drug dataset and the clinical labels. Further research is required to resolve those inconsistencies.

### Primacy of I_CaL_ and I_Kr_

The near-identical estimates of the axis of arrhythmia in *I*_CaL_ and *I*_Kr_ (Figure 5C) attest to the consistency between the biophysical model and the drug data for those two ion-currents, as well as the validity of using early after-depolarizations as a biomarker for drug-induced torsades de pointes. These consistencies are undoubtedly due to the close scrutiny of *I*_CaL_ and *I*_Kr_ over the past few decades. The primacy of *I*_CaL_ and *I*_Kr_ in predicting early after-depolarizations is consistent with the protective effect of co-expressed calcium and hERG potassium channels observed in ventricular cardiomyocytes (Ballouz et al., 2021). It is also consistent with previous findings in MICE (Multiple Ion Channel Effects) models (Kramer et al., 2013) where the torsadogenic risk of a drug is best predicted from the *IC*_50_ thresholds for *I*_CaL_ and *I*_Kr_ alone. MICE models are purely statistical and so have the same computational benefits as the axis of arrhythmia. However MICE models lack a biophysical interpretation, whereas the axis of arrhythmia describes the optimal changes in ion-channel conductivity (*G*_CaL_ and *G*_Kr_) to induce early after-depolarizations in the ventricular cardiac action potential.

### Inconsistency of I_Ks_

The biophysical model and the drug dataset produced conflicting estimates of the contribution of the slow potassium current (*I*_Ks_). The drug data returned a positive coefficient (*B*_Ks_ *>* 0) for the axis of arrhythmia (Figure 3C) which implies that *increasing G*_Ks_ is torsadogenic. Conversely, the biophysical model returned a negative coefficient (*B*_Ks_ *<* 0) which implies that *decreasing G*_Ks_ is torsadogenic. We suspect that the drug data is misleading in this case. Measurements of *I*_Ks_ vary greatly across experimental techniques (Ágoston et al., 2024) and only 58 of the 109 drugs in the present dataset had *IC*_50_ values for *I*_Ks_. Nonetheless, we cannot rule out errors in the biophysical model either. We concluded that *G*_Ks_ was an unreliable predictor of the present data, so we eliminated it from further consideration.

### Dilemma of I_NaL_

Similarly, inclusion of the late sodium current (*I*_NaL_) did not noticeably improve the predictive performance, so *G*_NaL_ was rejected as a poor predictor. However its exclusion did noticeably alter the risk scores for quinidine, thioridazine, quinine, ranolazine, lidocaine and mexiletine (Supplementary Information). Despite this, ranolazine was the only drug whose binary risk classification actually changed as a result. Using two ion-currents, ranolazine was erroneously classified as pro-arrhythmic (Figure 7B; solid line). Yet it was correctly classified with three ion-currents (Figure 7B; dotted line). This single outlier made little difference to the overall accuracy of the risk metric with two ion-currents (Figure 5D) versus three ion-currents (Figure 4D). We concluded that *I*_NaL_ is not a reliable predictor of torsadogenic risk in general. However, ranolazine remains the one exception whose predicted risk was rescued by the inclusion of *I*_NaL_. The metric in three ion-currents may therefore be informative for drugs that block *I*_Kr_ and *I*_NaL_ in a manner similar to ranolazine. Although, the present drug data suggests that it rarely improves the classification.

### Practical Risk Assessment

Our findings suggest that two ion-currents (*I*_CaL_ and *I*_Kr_) are sufficient to assess the pro-arrhythmic risk of novel drugs to an accuracy of between 89.9% and 91.7%. The fractional conductances (*δ*) of each ion-current are reconstructed from the *IC*_50_ measurements using Eq. (6). The pro-arrhythmic risk score is then computed from the action of the drug (*α* = ln *δ*) using the pooled coefficients of the axis of arrhythmia (Table 4). Specifically,

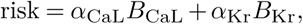

where *B*_CaL_ = 0.813 *±* 0.12 and *B*_Kr_ = *−*0.582 *±* 0.078. Positive risk scores indicate the drug is pro-arrhythmic, and negative risk scores indicate the drug is anti-arrhythmic. If the drug fails this test then it may be informative to compute the risk score using three ion-currents. Specifically,

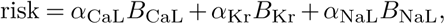

where *B*_CaL_ = 0.691 *±* 0.087, *B*_Kr_ = *−*0.617 *±* 0.069, and *B*_NaL_ = 0.377 *±* 0.075. The three-channel score may rescue the drug from a false-positive classification, as was the case for ranolazine. However, it should be noted that the biophysical model and the drug data disagree on the exact role of *I*_NaL_. So the predictions based on three ion-currents should be treated with caution.

### Closing Remarks

To the best our knowledge, the axis of arrhythmia is the only metric of pro-arrhythmic risk that can be derived from both (i) a biophysical model of the action potential, and (ii) a statistical model of drug-response profiles. It thus resolves the debate between biophysical and statistical models in safety pharmacology (Mistry, 2017; Lancaster and Sobie, 2017). More importantly, the existence of two methods allow independent lines of evidence to be compared. In this case, the biophysical model and the statistical model converged to near-identical estimates of the axis of arrhythmia in two ion-currents (*I*_CaL_ and *I*_Kr_). The available data revealed that the blockade of those two ion-currents was sufficient to predict the pro-arrhythmic risk of a drug with greater sensitivity and specificity than the standard hERG test with a 30-fold safety margin. The present metric is simple enough to be calculated with pen-and-paper, while retaining a meaningful biophysical interpretation. The score represents the propensity of the drug to induce early after-depolarizations in cardiomyocytes, averaged across cell phenotypes. Some phenotypes will be more vulnerable than others because they are naturally closer to the ectopic operating regime (Heitmann et al., 2023). The vulnerable proportion of the population can be calculated from the risk score, provided that the distributions of *G*_CaL_ and *G*_Kr_ are known (See Heitmann et al., 2023, for a discussion). Currently, those distributions are generally unknown. In terms of the current ICH S7B guidelines for pre-clinical risk assessment (ICH, 2005; Strauss et al., 2021), we believe that the inclusion of an *in vitro I*_CaL_ assay would augment the core *in vitro I*_Kr_ assay with minimal effort. Modern drug development pipelines would thus benefit from the improved accuracy of a multi-channel safety assay.

### Limitations

The drug data in the present study was originally collected from multiple laboratories under differing measurement protocols (Llopis-Lorente et al., 2020). Such measurements can differ substantially between recording sites, even when the same protocols are used (Kramer et al., 2020). In the present case, the ion-channel recordings for individual drugs were often amalgamated from multiple sources or had missing values. Of the 109 drugs in the dataset, only 17 drugs had no missing *IC*_50_ values for *I*_CaL_*, I*_Kr_*, I*_NaL_*, I*_Ks_. Likewise, only 21 drugs had no missing values for *I*_CaL_*, I*_Kr_*, I*_NaL_, and 83 drugs had no missing values for *I*_CaL_ and *I*_Kr_. Where values were missing, we assumed that the drug was known *a priori* not to block the ion-channels in question. The present study did not consider other potential triggers of arrhythmia, such as ectopic foci (Krummen et al., 2016) or fibrotic scar tissue (Jalife, 2000), because those phenomena are not induced by drugs.

## Methods

### Drug Dataset

The drug dataset is freely available from the source data of Heitmann et al. (2023). It was derived from the half-maximal inhibitory concentrations (*IC*_50_) and effective free therapeutic plasma concentrations (EFTPC) published in Supplementary Table S2 of Llopis-Lorente et al. (2020) which itself was curated from publicly available datasets and scientific publications. Where multiple values were encountered, Llopis-Lorente et al. (2020) used the median *IC*_50_ and the worst-case (highest) EFTPC values. The dose response curves were reconstructed from the *IC*_50_ values using the Hill equation (Hill, 1910),

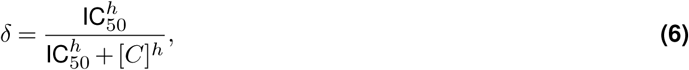

where *δ ∈* [0, 1] is the fractional conductance of the ion-channel. The concentration of the drug (*C*) was normalized to the effective free therapeutic plasma concentration (EFTPC). Ion-channels with missing IC_50_ values were assumed to be untouched by the drug (*δ* = 1). The Hill coefficients were assumed to be *h* = 1 in all cases. See Heitmann et al. (2023) for details. See Mirams et al. (2011) and Mirams (2023) for overviews.

### Clinical Risk Labels

The clinical risk labels for the drugs were transcribed from Table 1 of Llopis-Lorente et al. (2020), which was itself derived from the Credible Meds list of QT drugs (Woosley et al., 2019). The Credible Meds labeling scheme has four risk classes, where class 1 drugs carry a known torsadogenic risk; class 2 drugs carry a possible risk; class 3 drugs carry a risk but only in conjunction with external factors; class 4 drugs have no evidence of torsadogenic risk. The Credible Meds labels for each drug are included in the source data published by Heitmann et al. (2023). The present study used binary risk labels by lumping class 1 and 2 drugs together as ‘pro-arrhythmic’, and lumping class 3 and 4 drugs together as ‘benign’.

### Cross Validation

Leave-one-out cross-validation was used to train individual instances of the regression model (Eq. 4) on differing subsets of *n−*1 drugs, with one drug set aside for testing (n=109). The overall accuracy was calculated by averaging the test results across all *n −* 1 blocks. The resulting coefficients (*β*_ion_) and their standard errors (SE_ion_) were averaged across all blocks, after re-scaling to ensure that *||***B***||* = 1 in all cases.

### Pro-arrhythmic Risk Score

The risk score (Heitmann et al., 2023) was defined as,

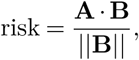

where **A** = *{α*_CaL_*, α*_Kr_*, α*_NaL_*, α*_Ks_*}* is the action of the drug of interest, and **B** = *{β*_CaL_*, β*_Kr_*, β*_NaL_*, β*_Ks_*}* is the basis vector of the axis of arrhythmia. The dot product,

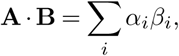

is the length of vector **A** after projecting it onto the axis of arrhythmia. The scores were normalized by scaling the basis vector **B** to unit length with the Euclidean norm,

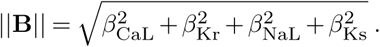

The basis vector points towards the pro-arrhythmic region of parameter space. Hence pro-arrhythmic drugs have positive risk scores, anti-arrhythmic drugs have negative risk scores, and neutral drugs have scores near zero.

### Scoring Threshold

Drugs were classified as *pro-arrhythmic* if their risk score exceeded a given threshold (risk *> θ*) and they were classified as *benign* otherwise (risk *< θ*). The optimal threshold at each dose (Figure 6B, red) was chosen to maximize the accuracy of the classifications, as described in Heitmann et al. (2023). The thresholds could also be adjusted to tune the balance of false-positives and true-positives in the ROC curve (Figure 6C).

### Biophysical Model

Following Heitmann et al. (2023), human ventricular action potential was simulated using a variant of the O’Hara et al. (2011) model that had been optimized to reproduce the long QT phenotype (Krogh-Madsen et al., 2017). Randomization of the ionic conductances was limited to those ion-channels that served as predictor variables in the regression analysis. Namely, (*G*_CaL_, *G*_Kr_, *G*_NaL_) *∈* [*e^−^*^3^*, e*^+3^] or (*G*_CaL_, *G*_Kr_) *∈* [*e^−^*^3^*, e*^+3^]. In both cases the sample size was *n*=10000 cardiomyocytes. Those myocytes that elicited early after-depolarization were labeled as *ectopic* and those that did not were labeled as *benign*. Those labels formed the basis of the logistic regression analysis of the cell model.

### Pooled Estimate of the Axis

The coefficients of the competing axes of arrhythmia (Figure 6) were pooled by simple average,

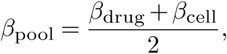

and re-scaled to preserve *||***B***||* = 1. The pooled standard errors,

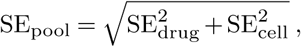

were re-scaled too. The confidence intervals of the risk scores (Figure 7) were computed from these standard errors.

### Statistics

Welch’s two-sample t-statistic,

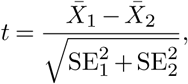

was used to compare regression coefficients having unequal variances and sample sizes. The degrees of freedom were calculated as *df* = *a/*(*b* + *c*) where 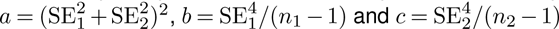.

### Source Code Availability

The source code for the biophysical model is available from https://zenodo.org/records/7796721 under the GNU General Public License v3.0. The cell model requires version 2022 or later of the Brain Dynamics Toolbox (Heitmann et al., 2018; Heitmann and Breakspear, 2022) which can be downloaded from https://zenodo.org/records/7070703 under the BSD 2-clause license.

## Acknowledgements

This study was supported by Australian NHMRC grants app1182032, app1182623 and NSW Health cardiovascular capacity building grant. SH was funded by The Medical Advances Without Animals Trust (MAWA) which aims to advance medical science and improve human health and therapeutic interventions without the use of animals or animal products. The Katana computing cluster on which the simulations were conducted is supported by Research Technology Services at UNSW Sydney.

## Supplementary Information

The multi-channel (*I*_CaL_, *I*_Kr_, *I*_NaL_, *I*_Ks_) blocking profiles of each drug in the dataset, grouped by the torsadogenic risk labels from Credible Meds. In that scheme: **Class 1** drugs (red) carry a known torsadogenic risk; **Class 2** drugs (dark orange) possibly carry a risk; **Class 3** drugs (light orange) carry a risk, but only in conjuction with external factors; **Class 4** drugs (yellow) have no evidence of torsadogenic risk. The source data for the drugs can be downloaded from Table 2 in Heitmann, et al. (2023). The trajectories of multi-channel block are plotted below. The pro-arrhythmic risk scores were computed using the pooled axis of arrhythmia in two ion-currents (heavy black line) and also in three ion-currents (dotted line) as described in the main text.

### Drugs with Class 1 Torsadogenic risk

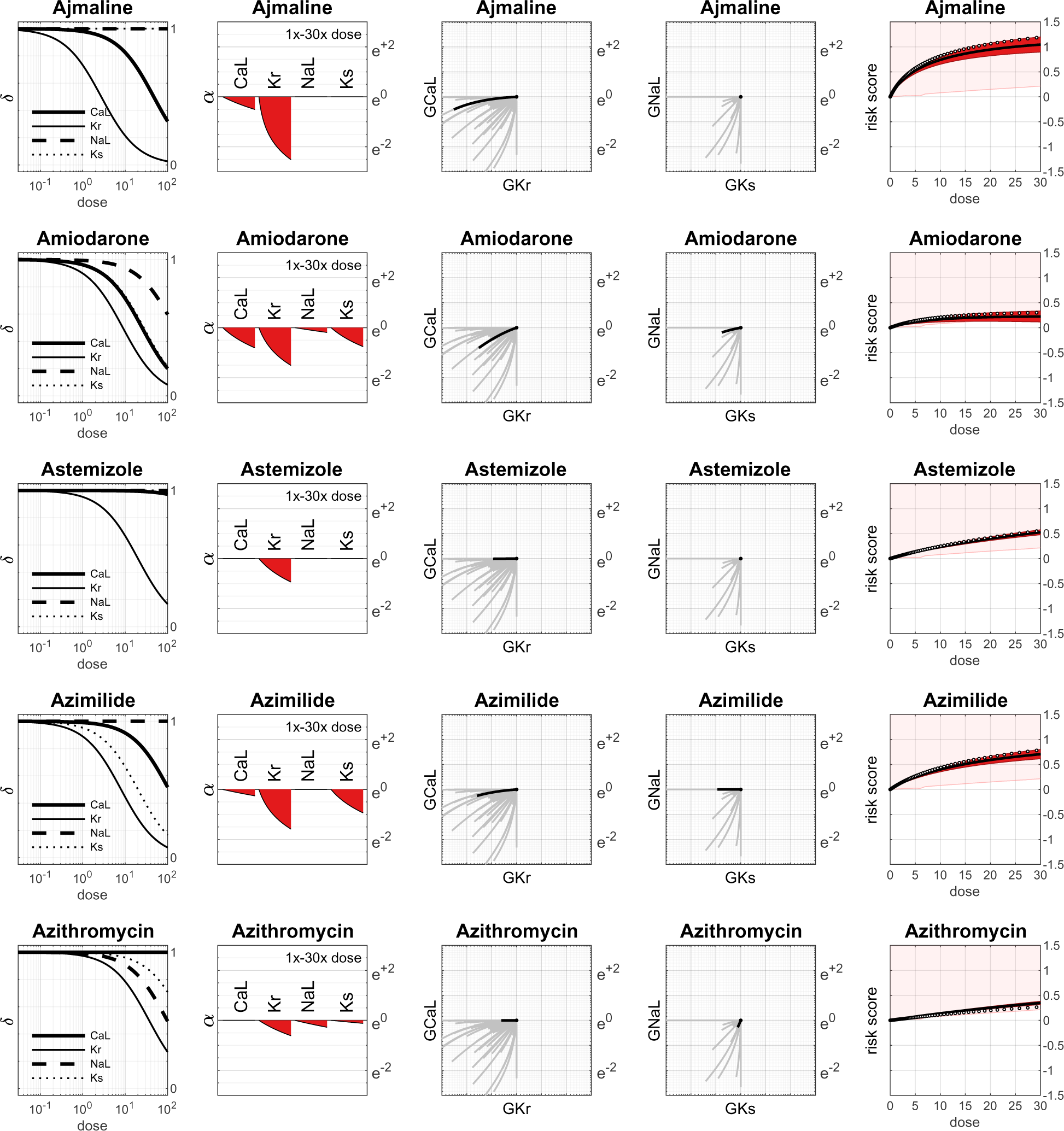
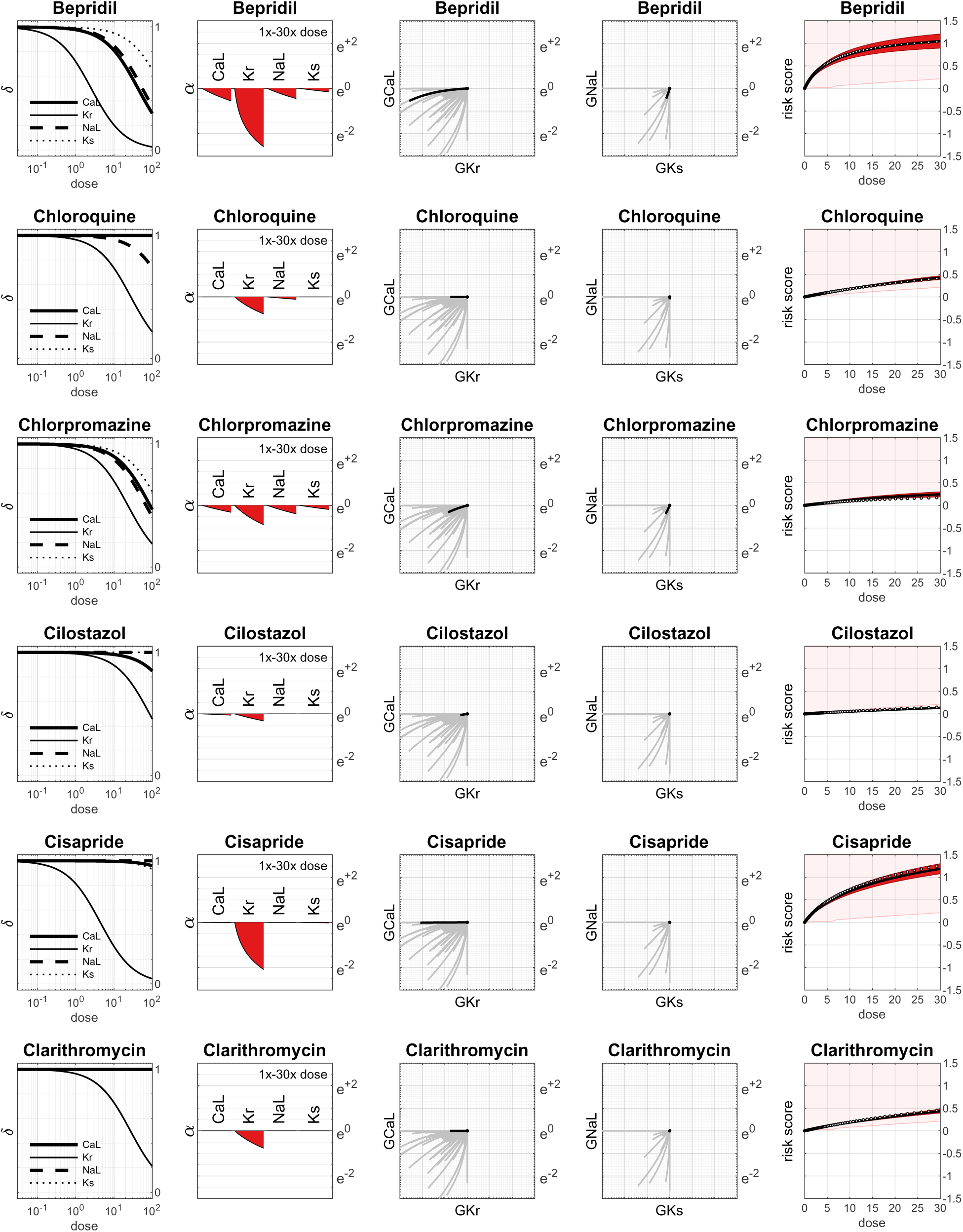
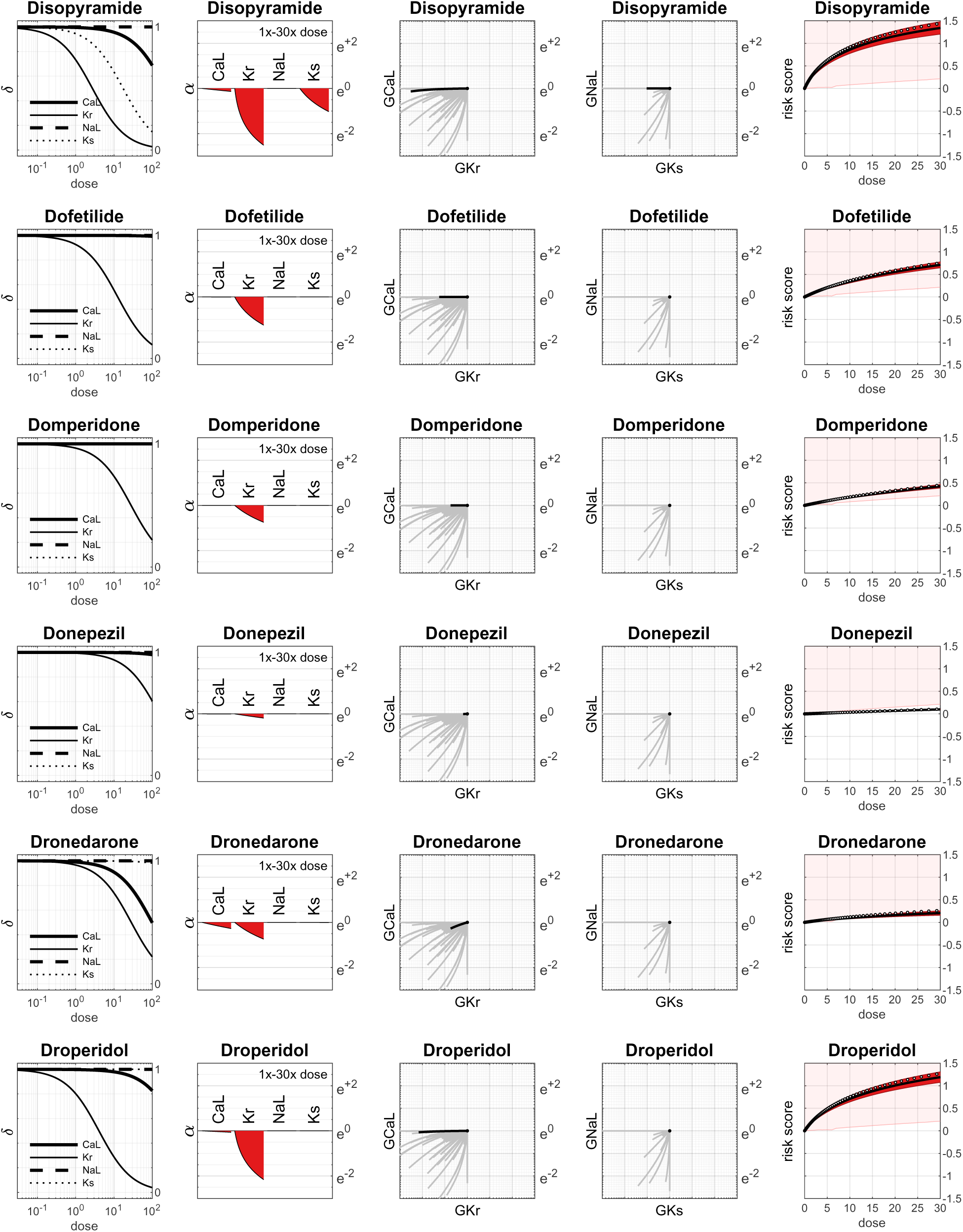
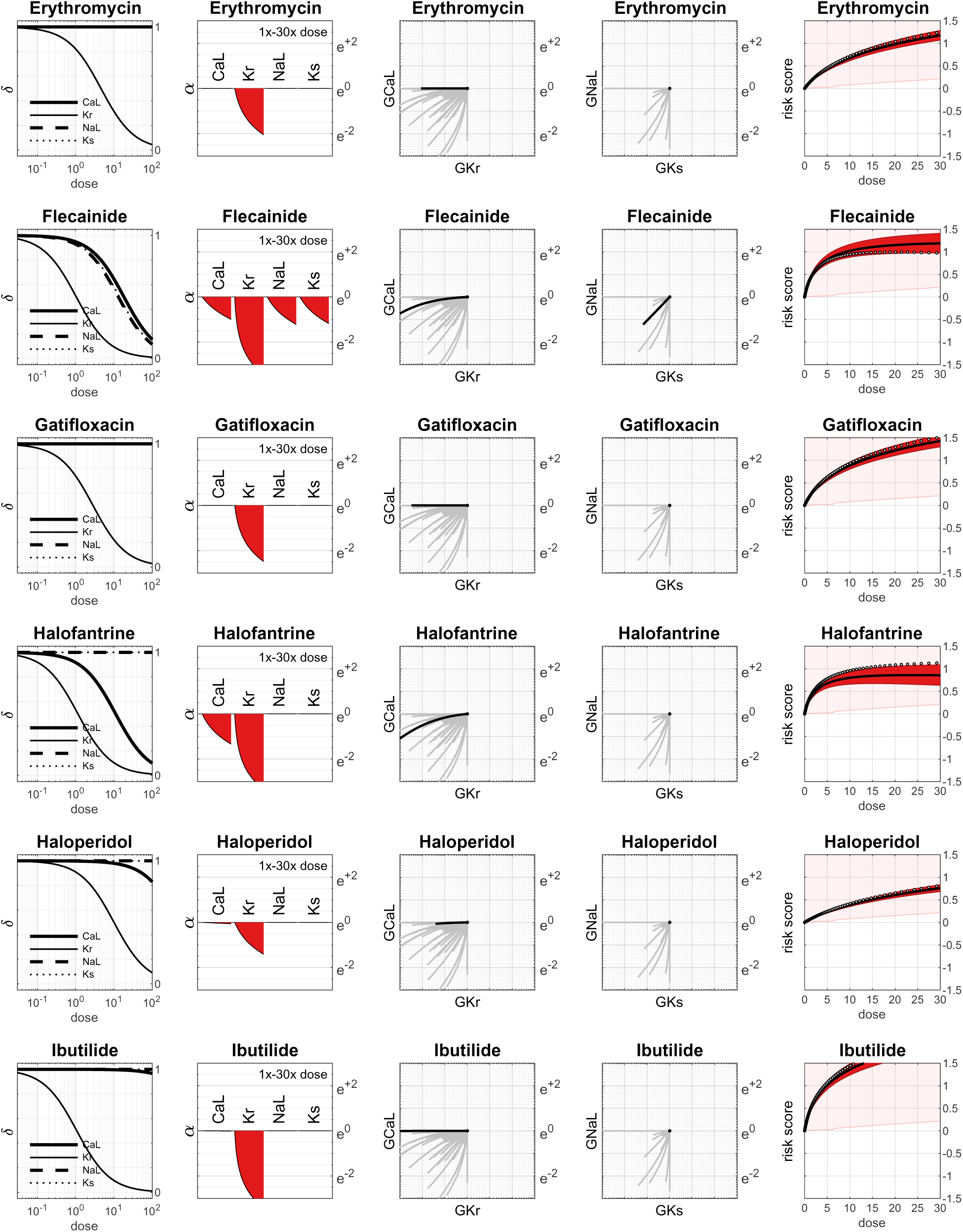
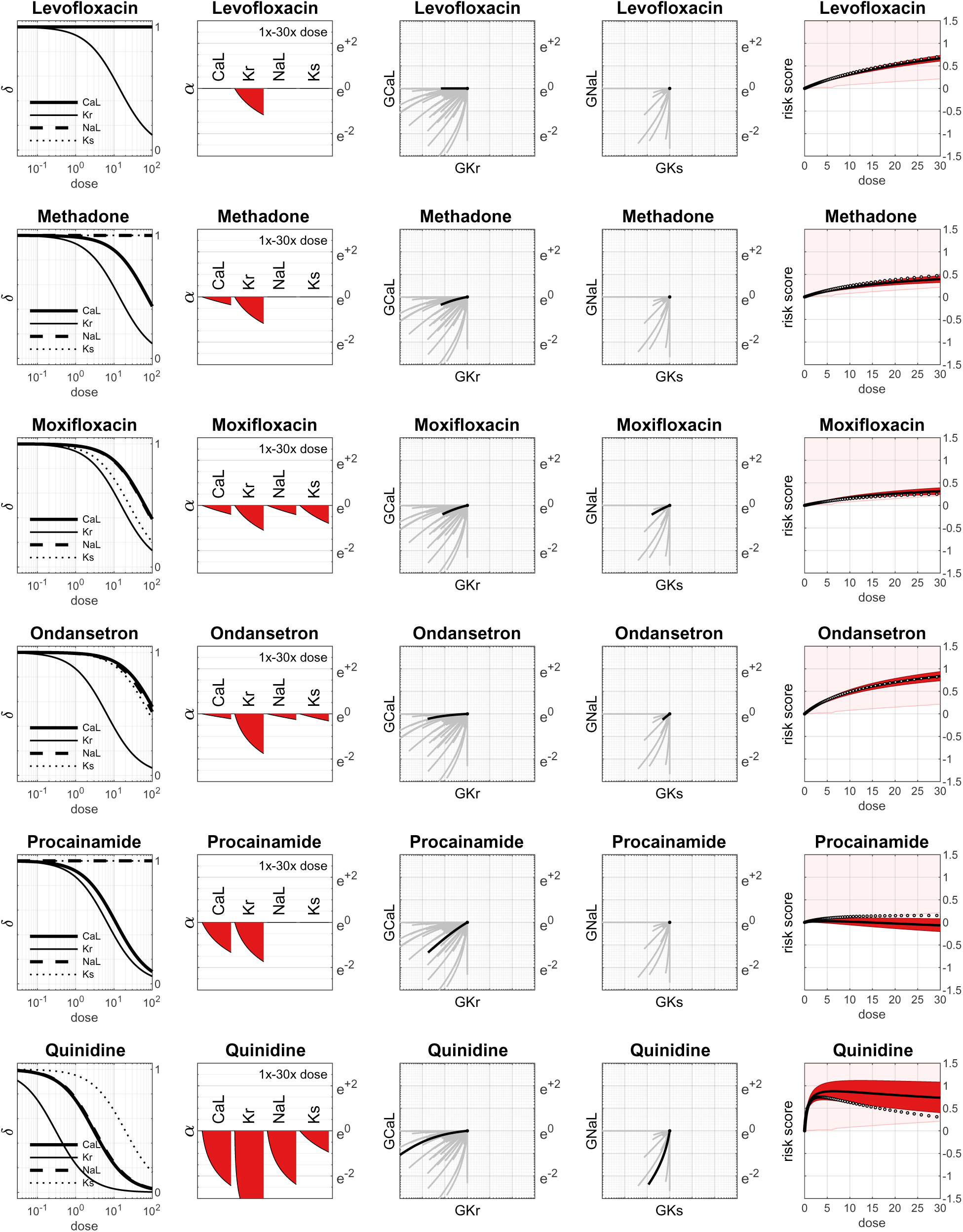
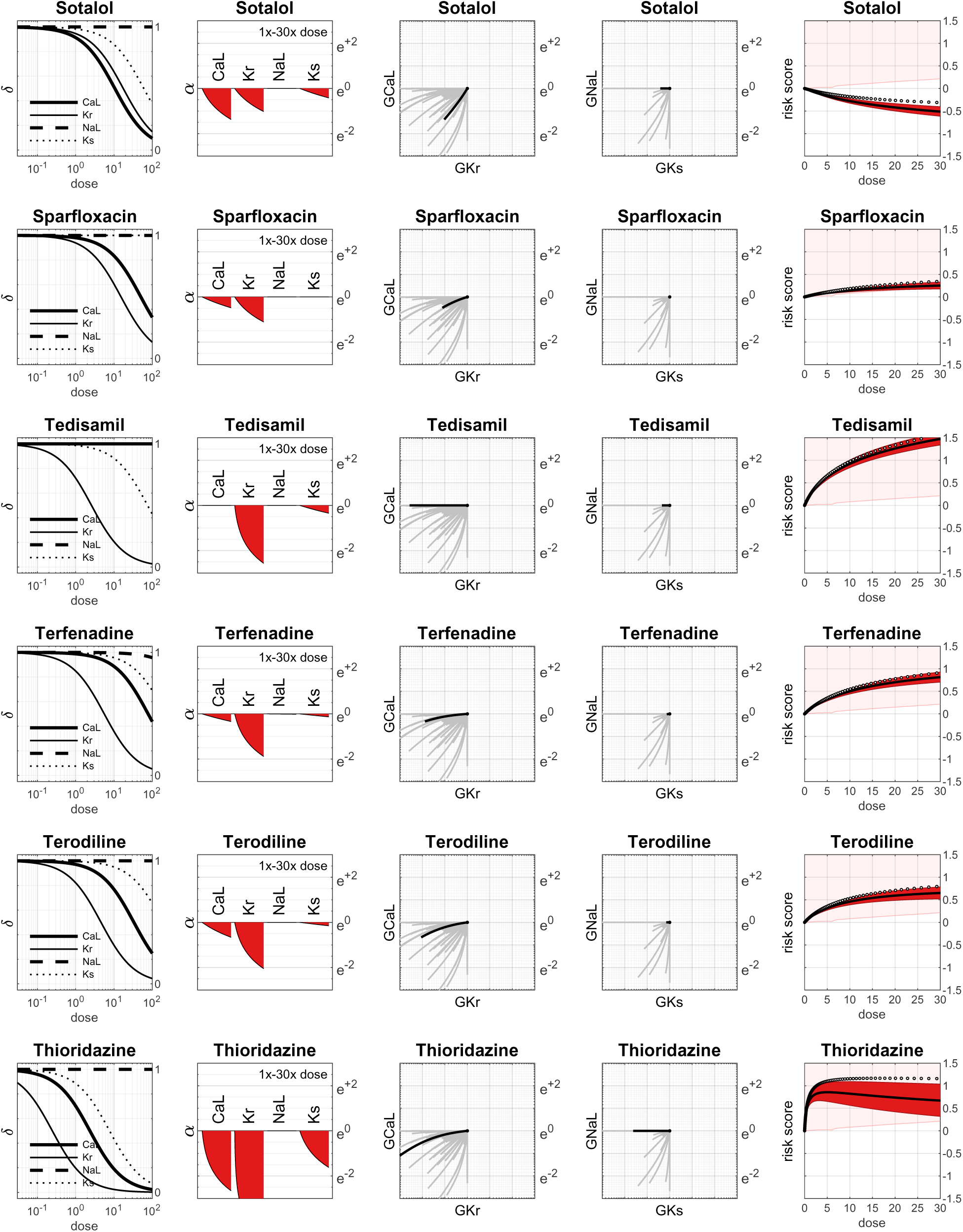
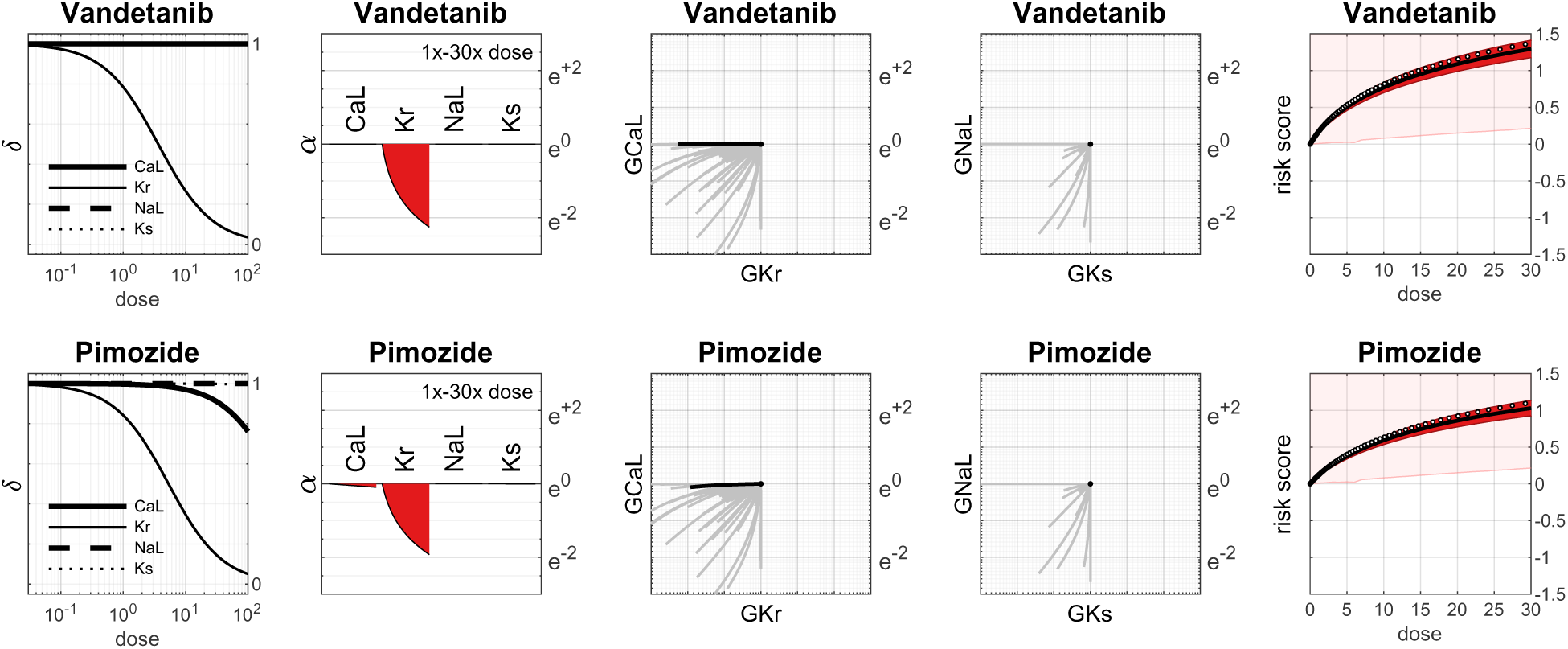

### Drugs with Class 2 Torsadogenic risk

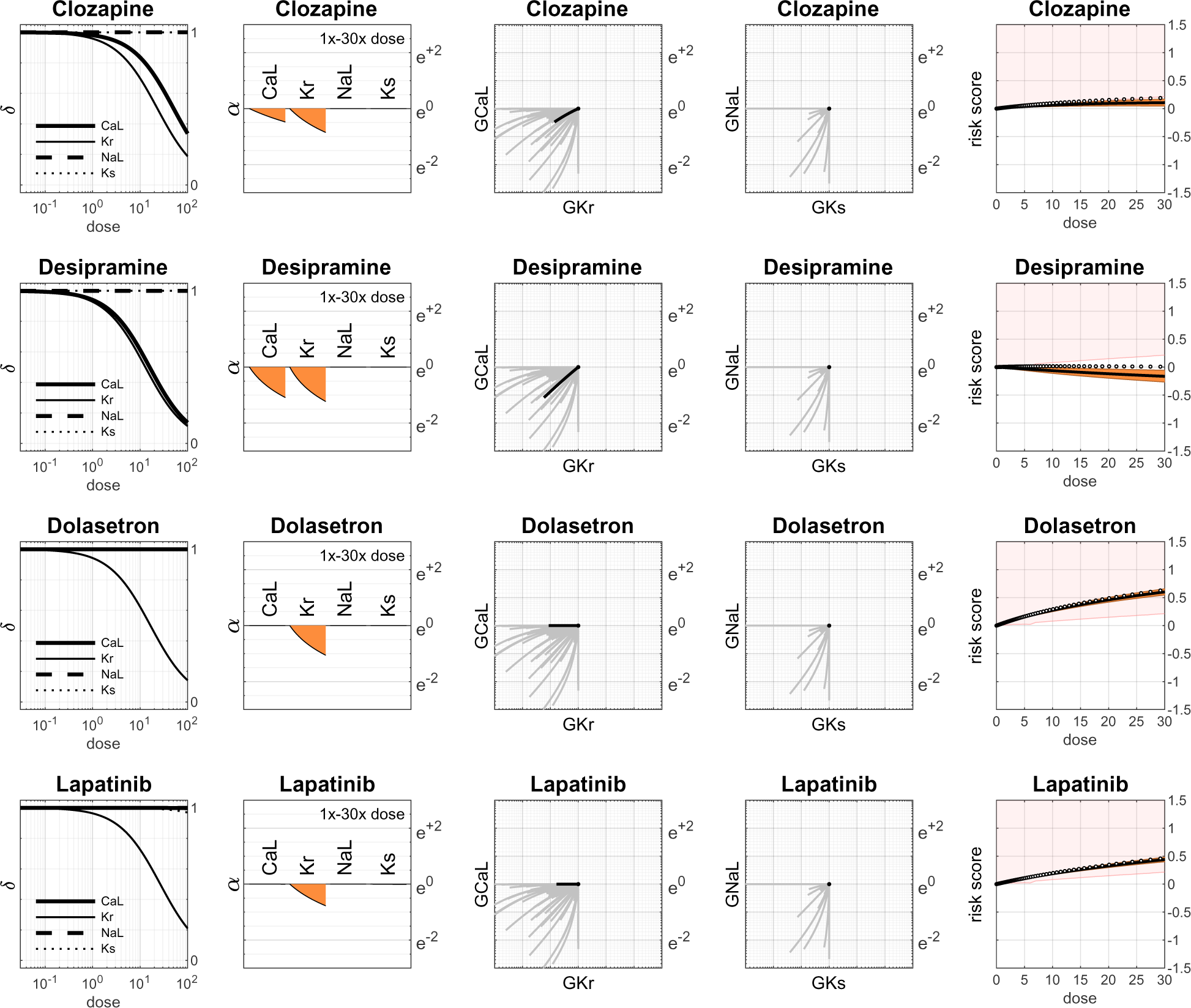
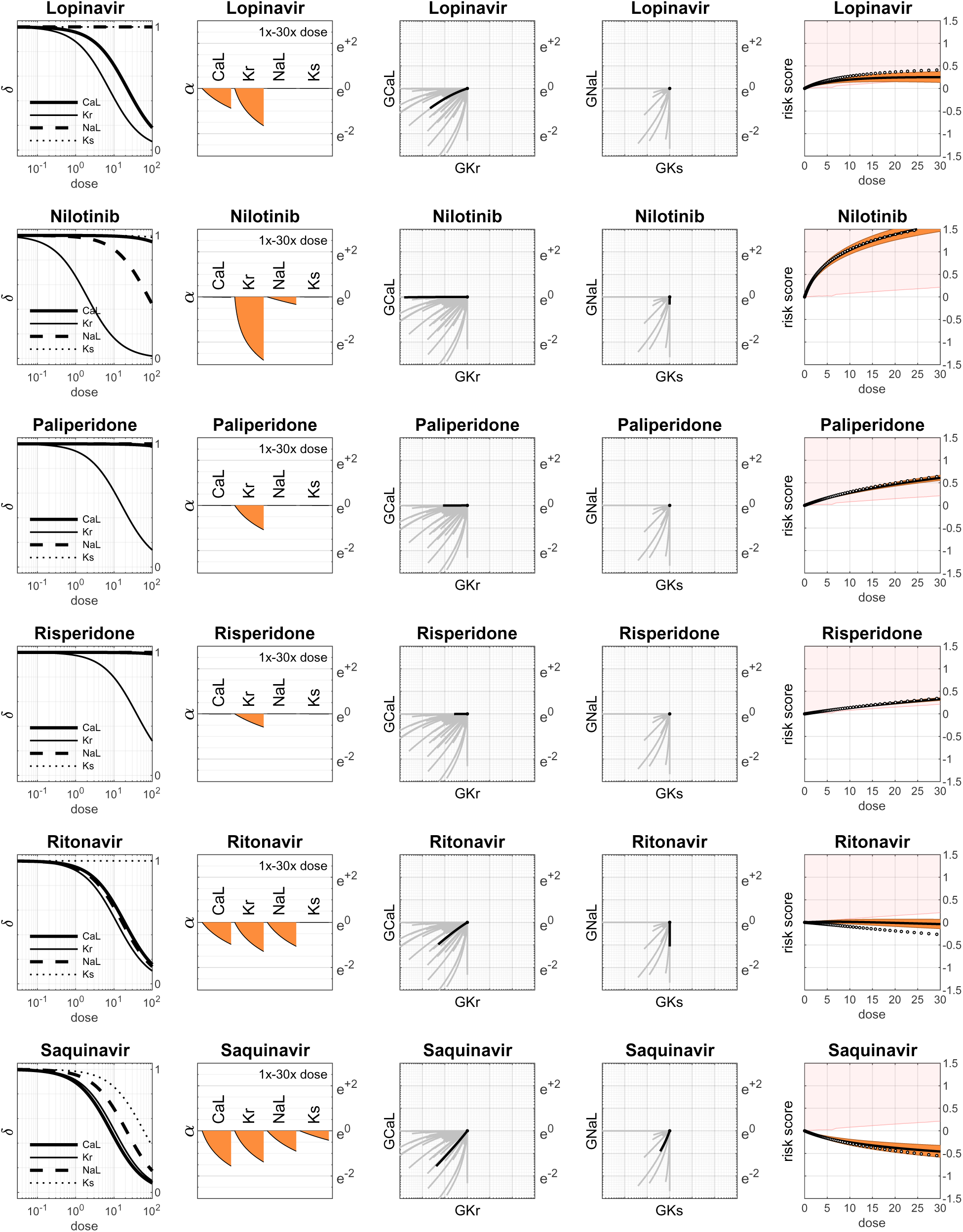
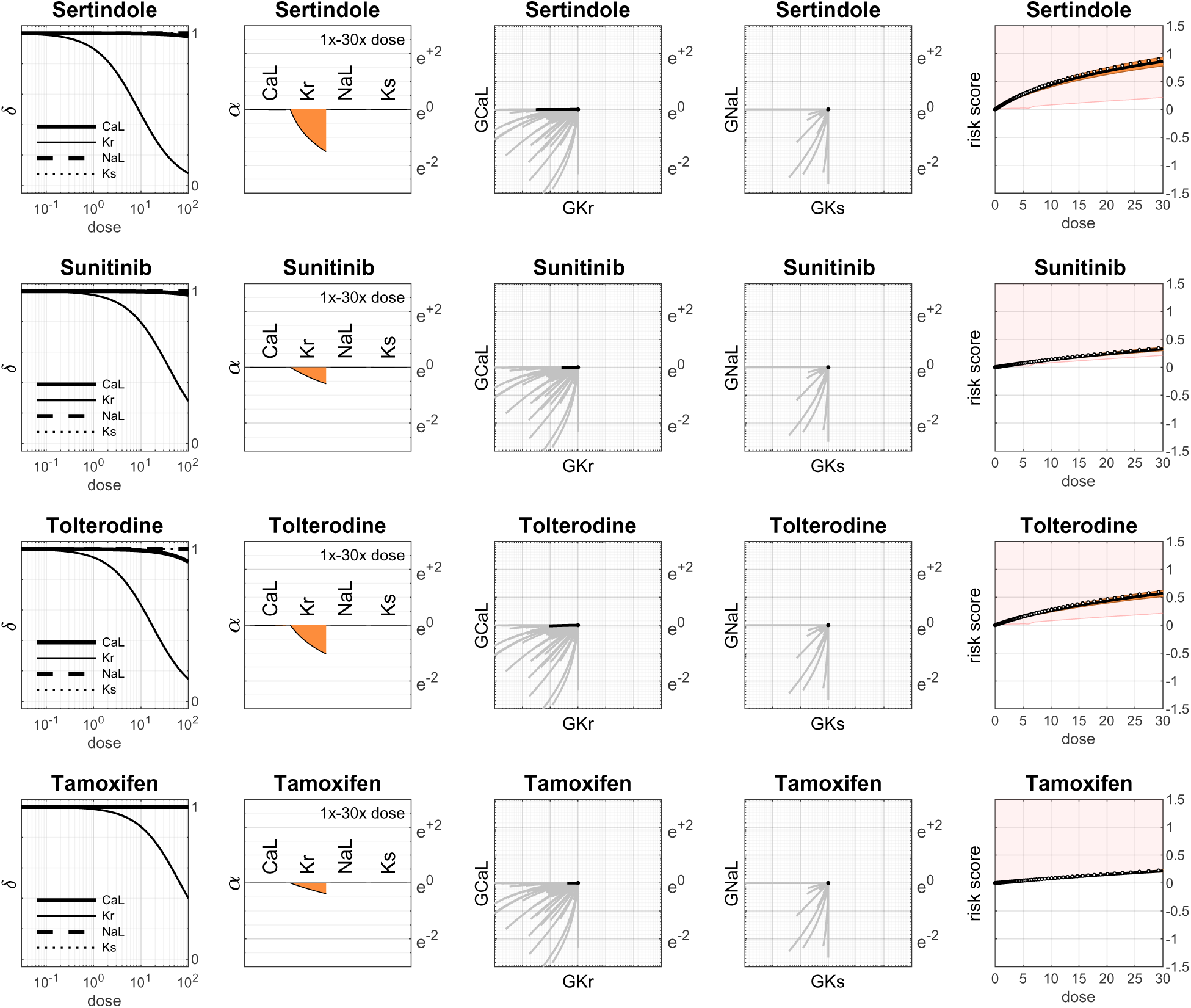

### Drugs with Class 3 Torsadogenic risk

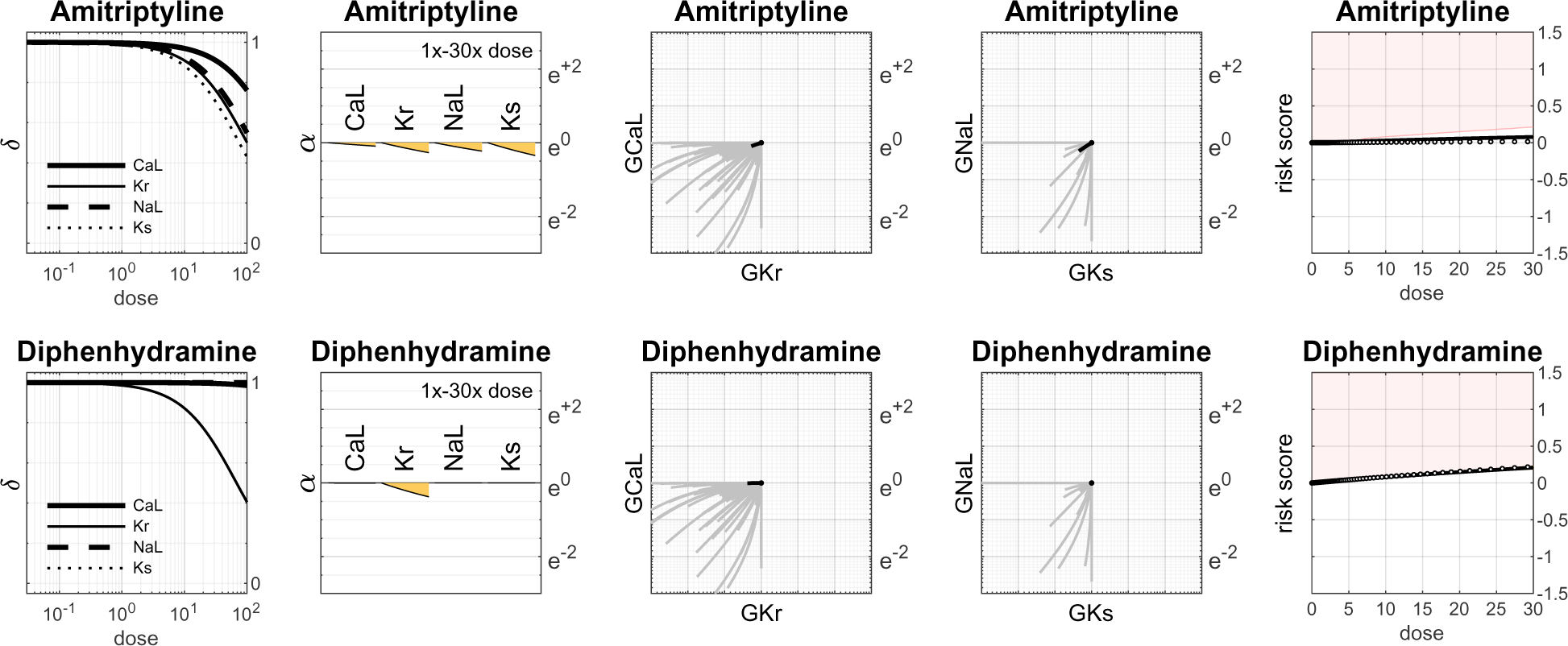
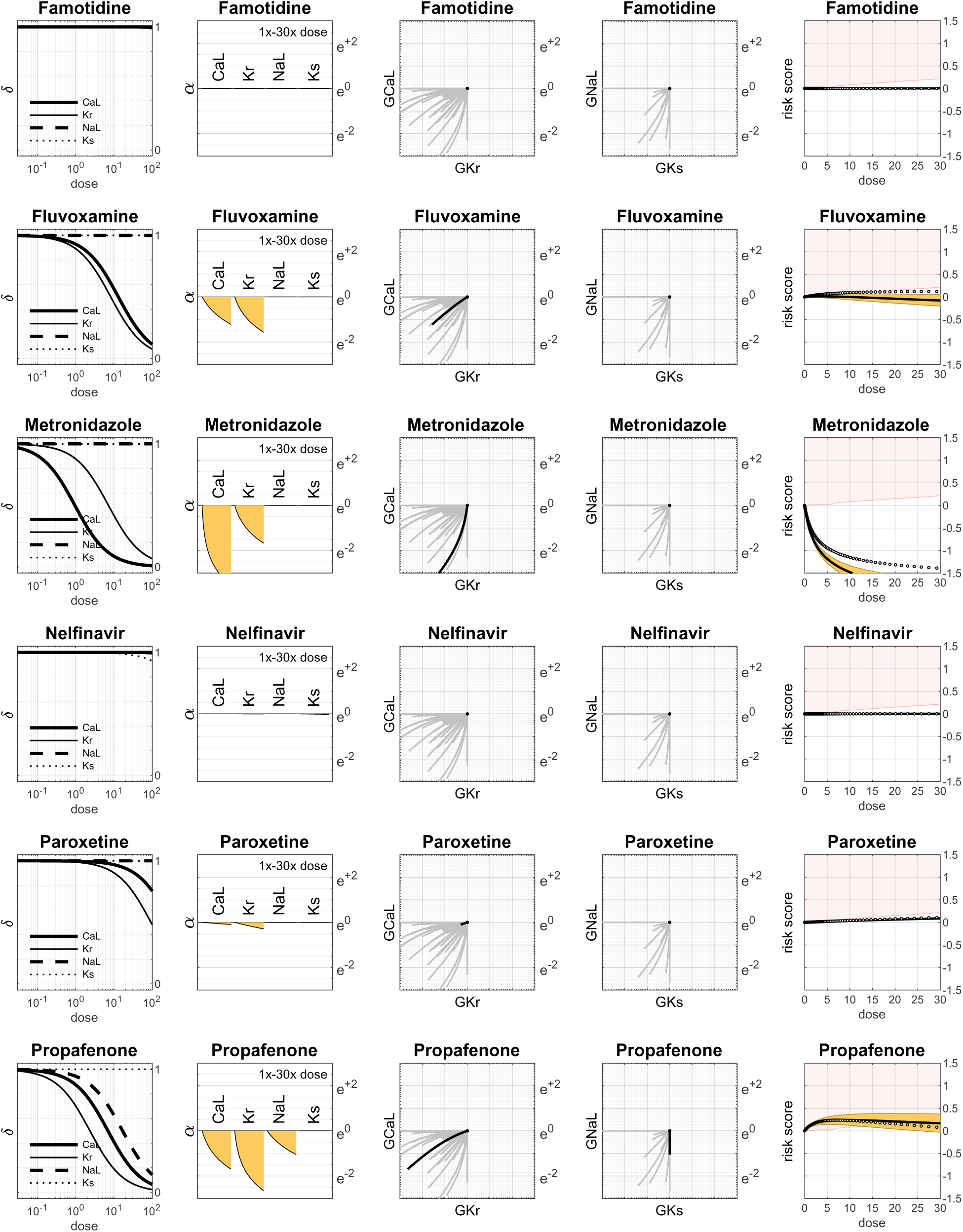
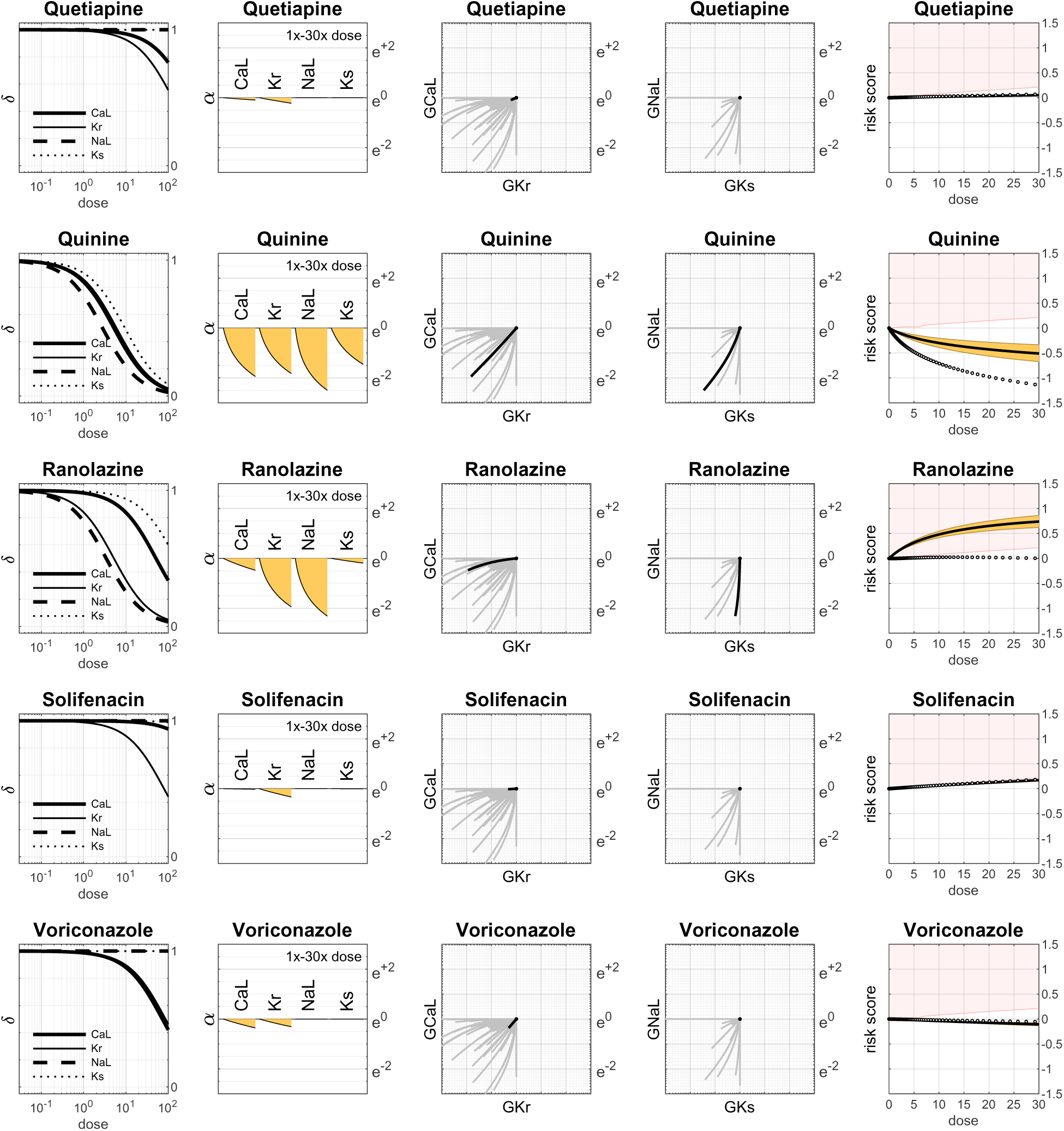

### Drugs with Class 4 Torsadogenic risk

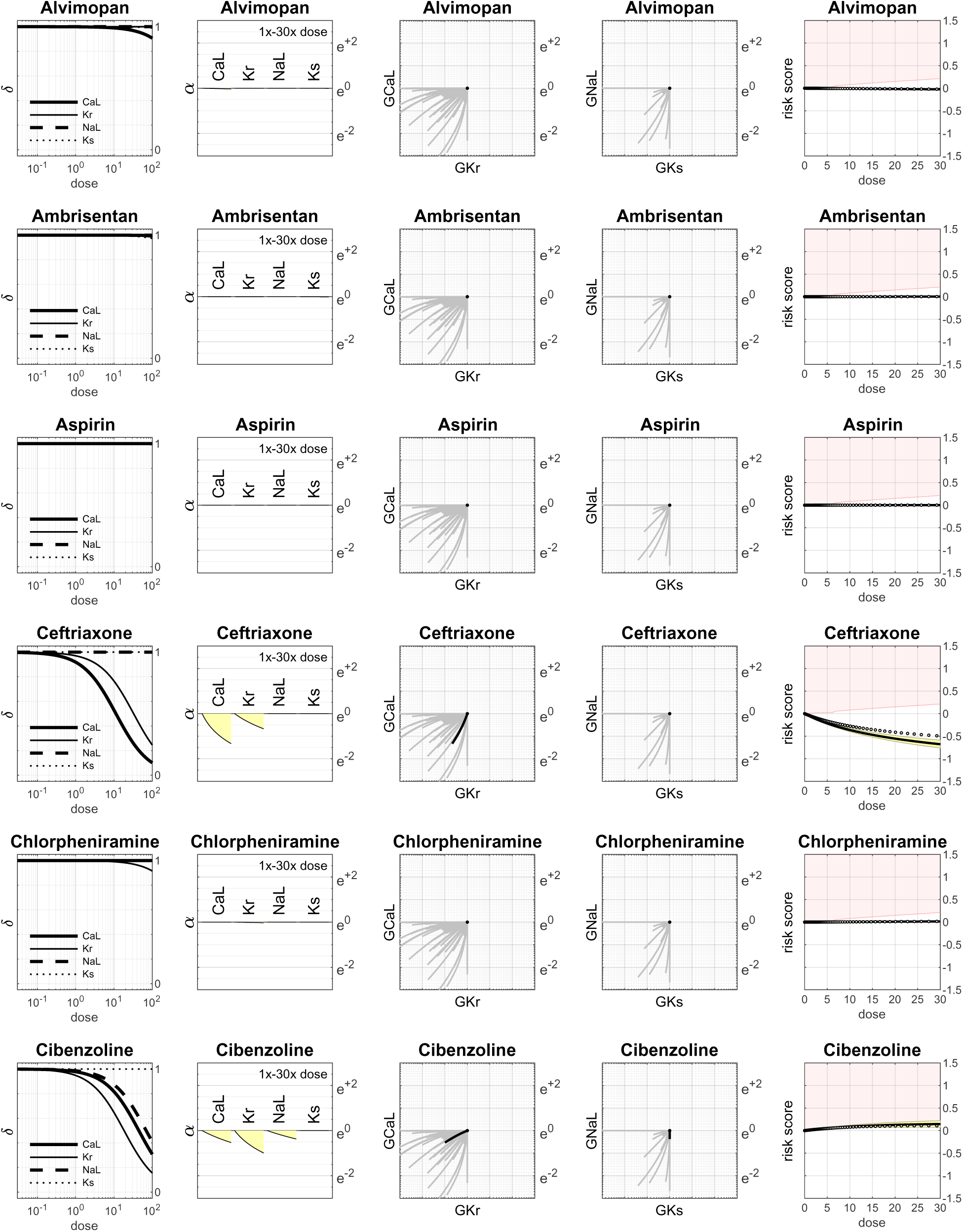
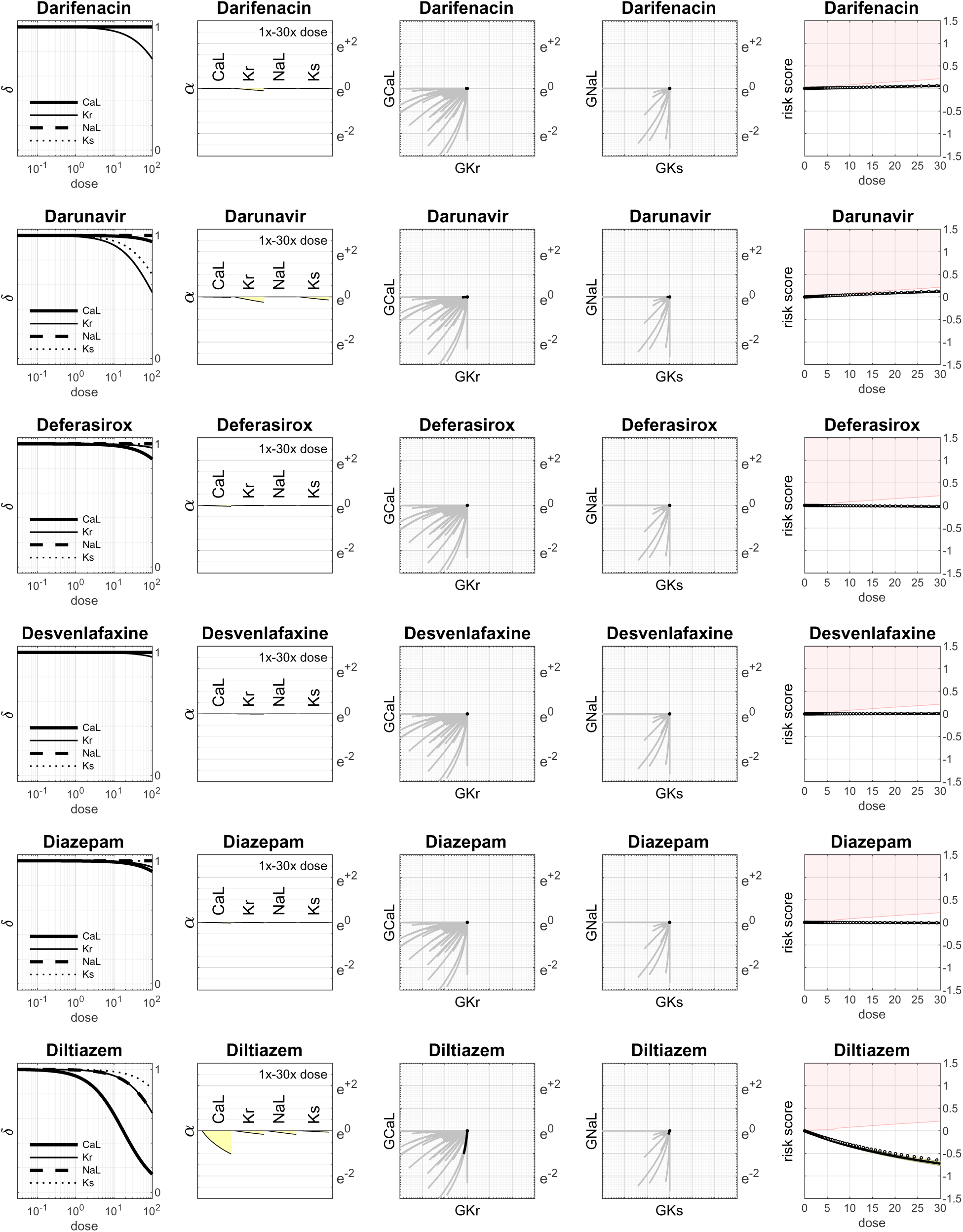
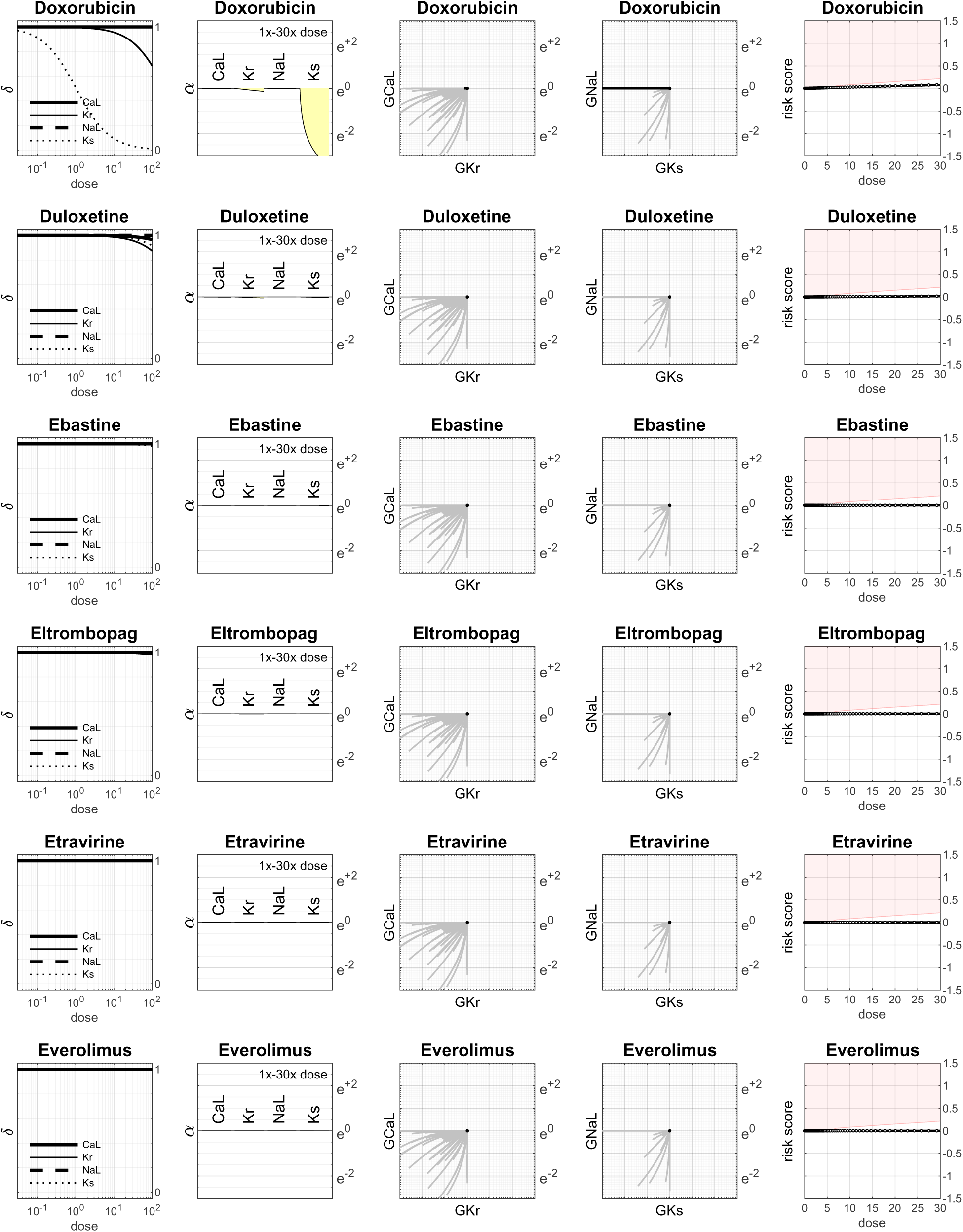
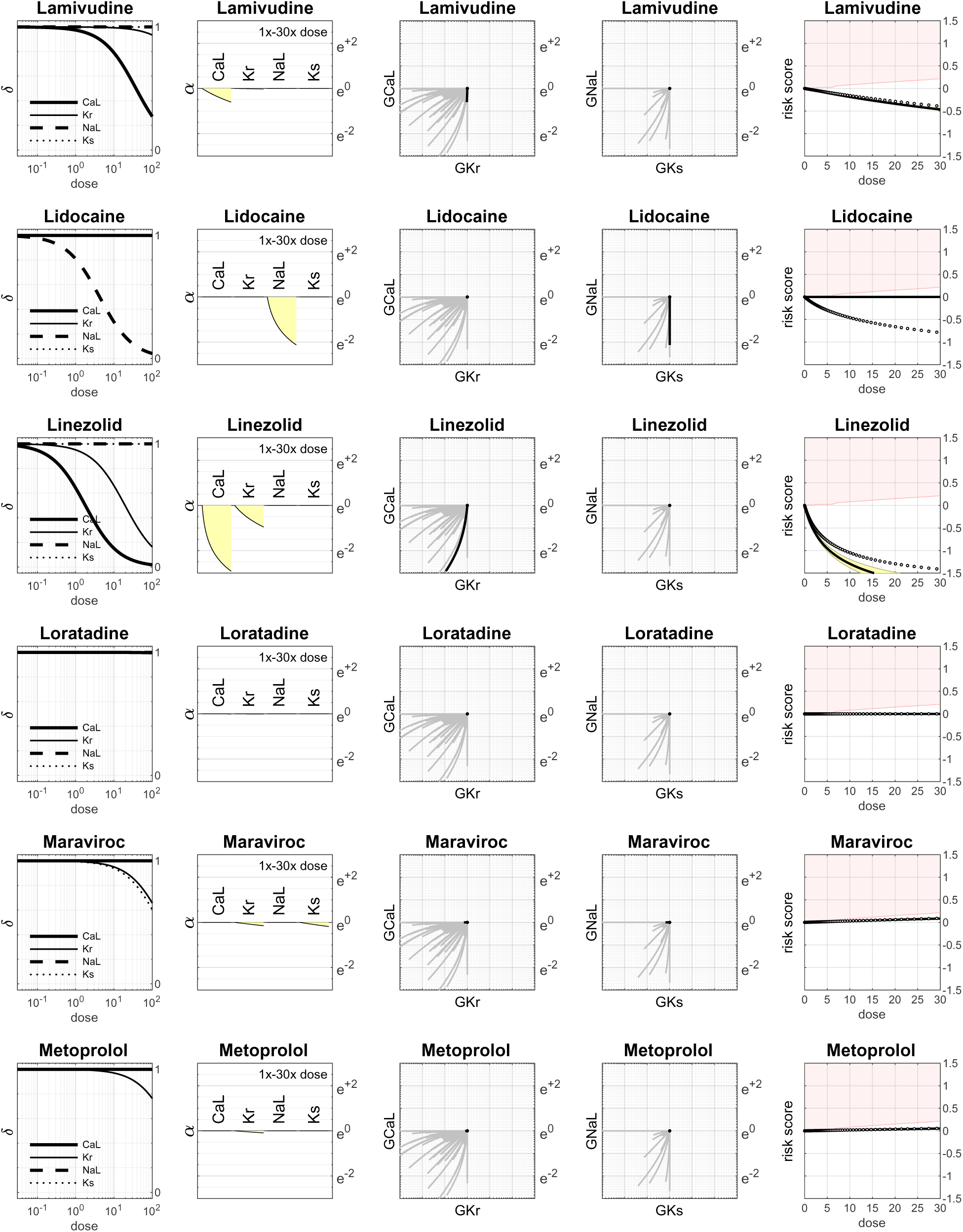
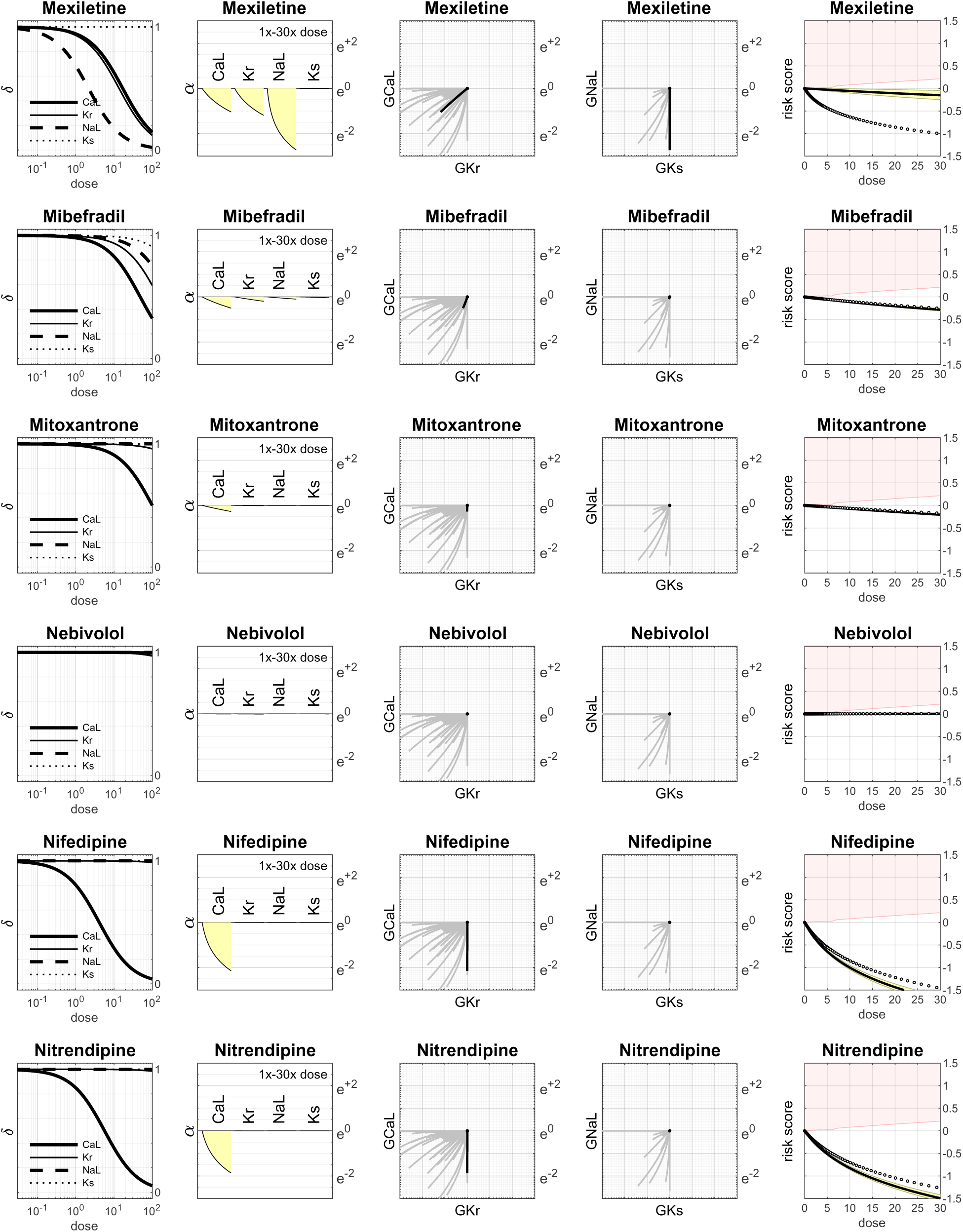
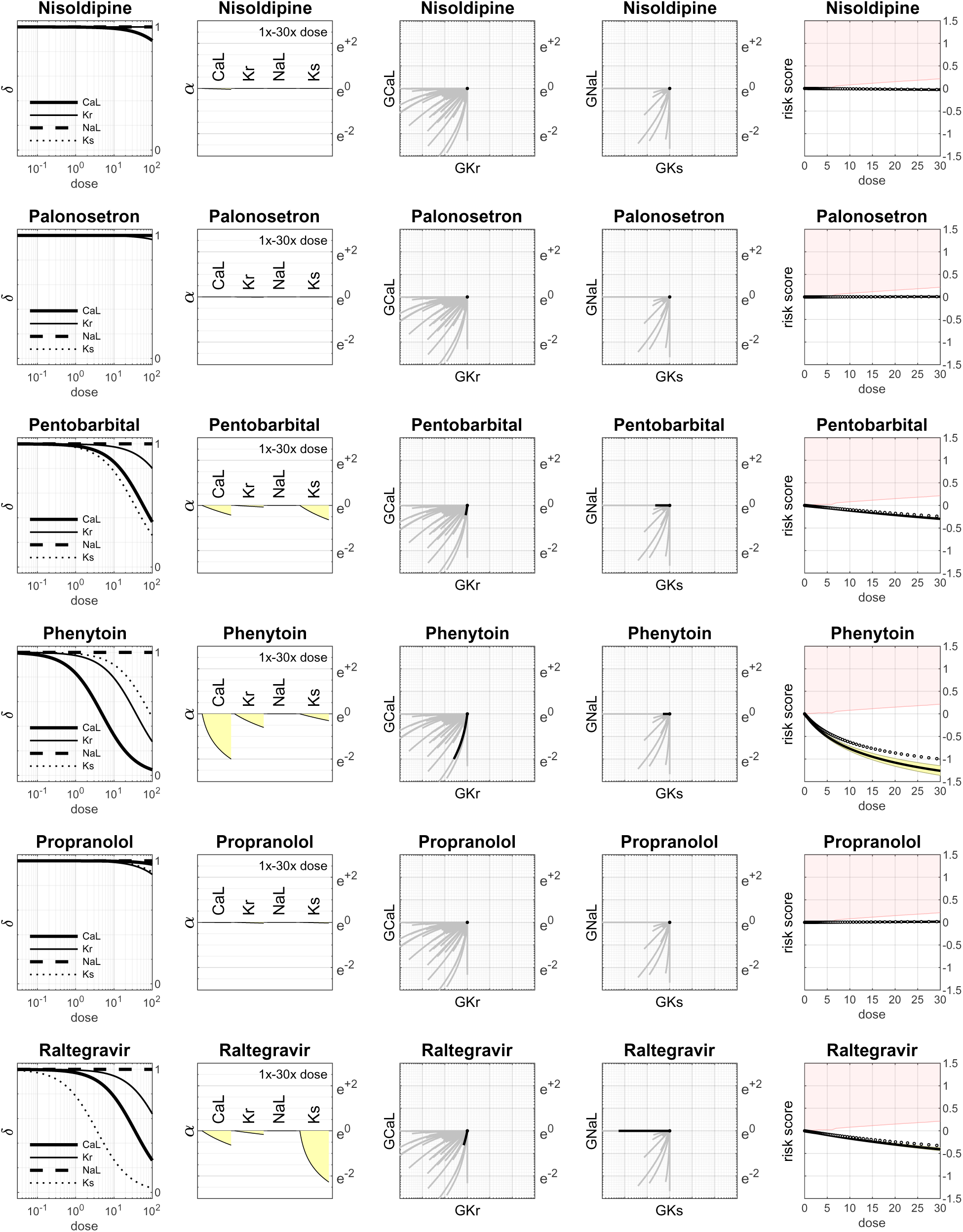
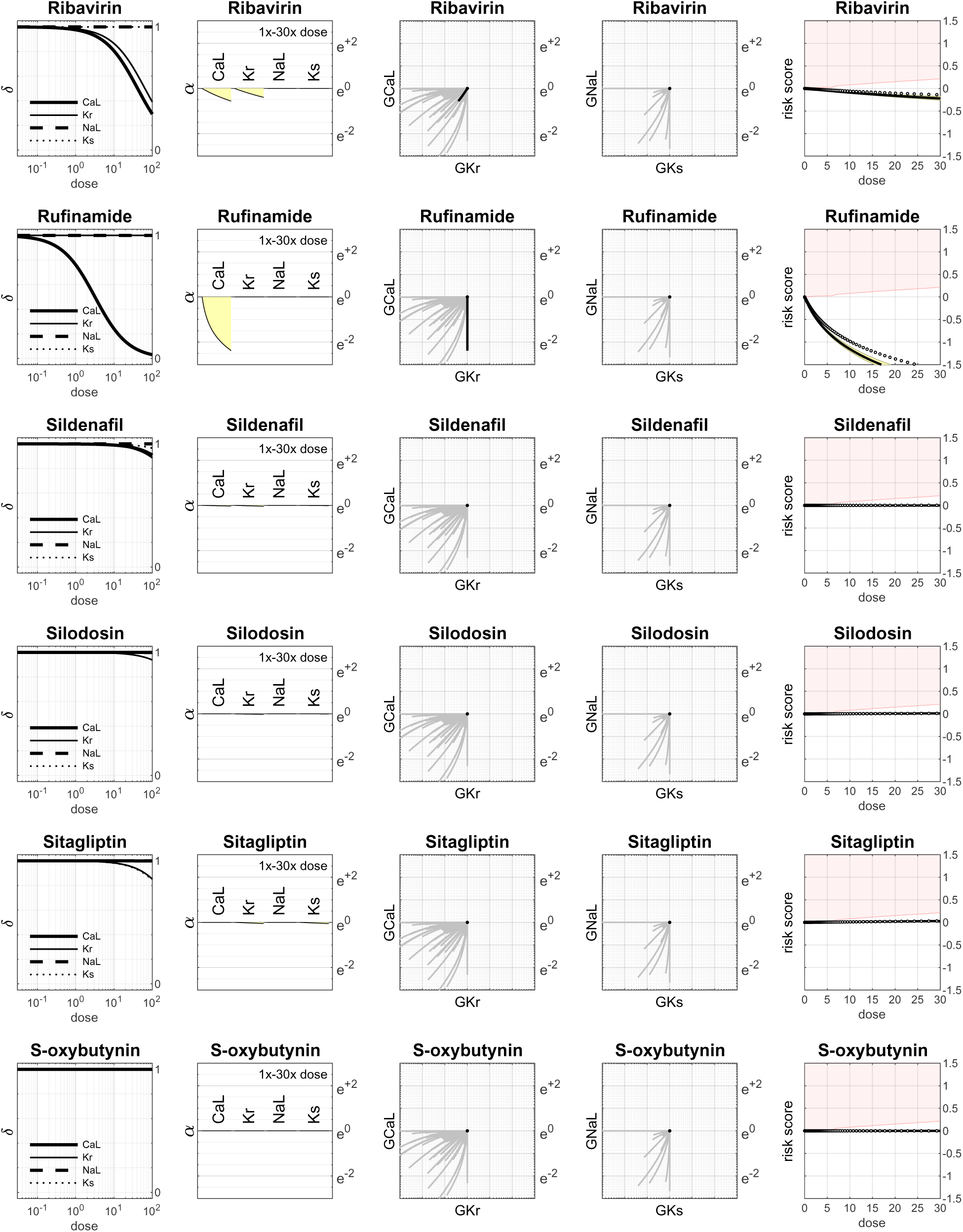
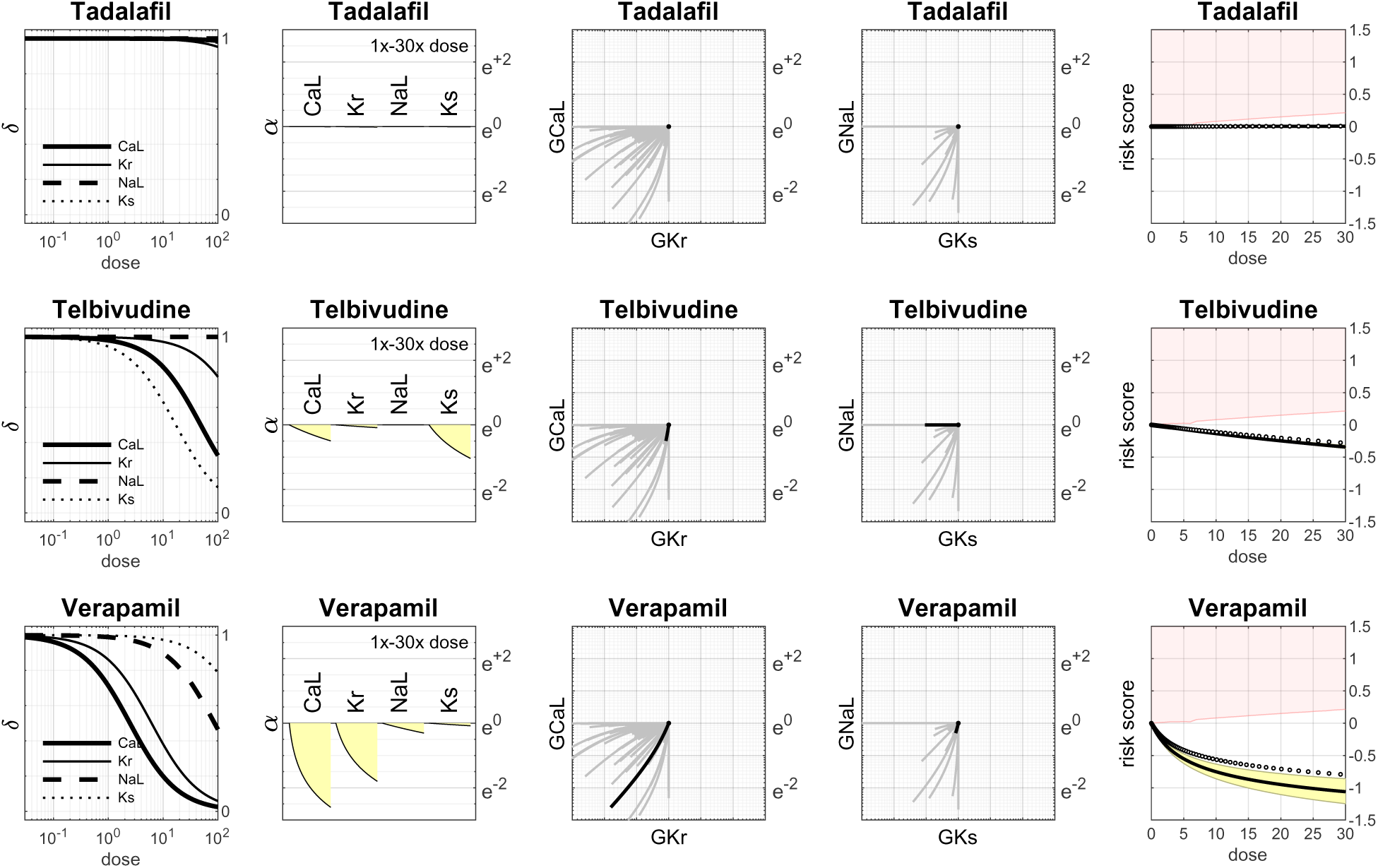

